# Interplay of two small RNAs fine-tunes hierarchical flagellar gene expression in the foodborne pathogen *Campylobacter jejuni*

**DOI:** 10.1101/2023.04.21.537696

**Authors:** Fabian König, Sarah L. Svensson, Cynthia M. Sharma

**Affiliations:** University of Würzburg, Institute of Molecular Infection Biology, Department of Molecular Infection Biology II, 97080 Würzburg, Germany; The Center for Microbes, Development and Health, CAS Key Laboratory of Molecular Virology and Immunology, Institut Pasteur of Shanghai, Chinese Academy of Sciences, Shanghai, China 200031

**Keywords:** *Campylobacter jejuni*, small RNAs, RNase III, RNase Y, PNPase, post-transcriptional regulation, motility, flagellar biogenesis

## Abstract

Like for many enteric bacteria, flagella are a crucial virulence factor for the foodborne pathogen *Campylobacter jejuni*, allowing the bacteria to move through the viscous mucus of the human intestine. Assembly of the complex flagellar machinery and filament requires hierarchical regulation via transcriptional control of each component. In *C. jejuni*, class I genes are transcribed from σ^70^-dependent promoters and class II/III genes with the help of the alternative sigma factors RpoN (σ^54^) and FliA (σ^28^). In contrast to transcriptional control, less is known about post-transcriptional regulation of flagellar biosynthesis cascades via small regulatory RNAs (sRNAs). Here, we characterized two sRNAs with opposing effects on the cascade that fine-tune *C. jejuni* flagellar filament assembly and thereby impact motility. We demonstrate that the highly conserved *Campylobacter* sRNA CJnc230 (FlmE, flagellar length and <u>m</u>otility <u>e</u>nhancer), encoded downstream of the flagellar hook structural protein FlgE, is dependent on RpoN and that RNase III processes CJnc230 from the *flgE* mRNA, while RNase Y and PNPase mature its 3’ end. We identify mRNAs encoding a regulator of flagella-flagella interactions and the anti-σ^28^ factor FlgM as direct targets of CJnc230 repression. Overexpression of CJnc230 de-represses FliA activity and upregulates class III flagellar genes, such as the major flagellin *flaA*, culminating in longer flagella and increased motility. In contrast, overexpression of the FliA-dependent sRNA CJnc170 (FlmR, flagellar length and <u>m</u>otility <u>r</u>epressor) reduces flagellar length and motility. Overall, our study demonstrates sRNA-mediated post-transcriptional regulation fine-tunes *C. jejuni* flagellar biosynthesis through balancing of the hierarchically expressed components.

## Introduction

Flagella are crucial for colonization of the human gut by many bacterial pathogens. They mediate not only chemotaxis towards the epithelial lining, but also adhesion to host cells, immunogenicity, and even secretion of virulence factors (Akahoshi and Bevins, 2022; Colin et al., 2021; Haiko and Westerlund-Wikström, 2013; Scanlan et al., 2017). Their biogenesis in gammaproteobacterial model organisms such as *Escherichia coli* and *Salmonella enterica* follows a well characterized cascade of hierarchical transcriptional control to mediate assembly of the flagellum from the inside out (Chevance and Hughes, 2008). In *E. coli* and *Salmonella*, early (class I) flagellar genes include the master transcriptional regulator FlhDC, which activates middle (class II) genes, mainly comprising hook-basal-body (HBB) complex components, as well as the sigma factor σ^28^ (FliA) and its antagonist FlgM (Gillen and Hughes, 1991). Upon completion of the HBB, FlgM is secreted, thereby releasing FliA for transcriptional activation of late flagellar genes (class III), including the filament component flagellin (Chevance and Hughes, 2008). Despite flagella being a paradigm for studying hierarchical gene expression in bacteria, their regulation varies among different species and little is known about post-transcriptional control in this complex process.

The Epsilonproteobacterium *Campylobacter jejuni* is the currently most prevalent cause of bacterial foodborne illness worldwide (Havelaar et al., 2015). While its small ∼1.6 megabase pair (Mbp) genome lacks classical virulence factors encoded by enteric pathogens, such as dedicated type III secretion systems (Parkhill et al., 2000), *C. jejuni* secretes virulence-associated factors through its flagellar type III secretion system (fT3SS) (Burnham and Hendrixson, 2018). Moreover, flagella-mediated motility is also a crucial *C. jejuni* virulence determinant (Gao et al., 2017; Guerry, 2007; Liu et al., 2012), and the flagellum mediates host cell adhesion (McSweegan and Walker, 1986). In contrast to *E. coli* or *Salmonella*, Epsilonproteobacteria do not encode a homolog of the master regulator FlhDC, and early genes in the hierarchical transcriptional cascade are expressed constitutively with the help of the housekeeping sigma factor RpoD (σ^70^ in *C. jejuni*) (Lertsethtakarn et al., 2011). Moreover, Epsilonproteobacteria have dedicated their two additional sigma factors (RpoN and FliA) to drive expression of middle and late flagellar genes (**Fig. 1A**) (Gilbreath et al., 2011). In *C. jejuni*, early genes comprise components of the fT3SS, the cytoplasmic and inner membrane rings, as well as activators of class II gene expression, including the two-component system factors FlgS (histidine kinase) and FlgR (σ^54^-associated transcriptional activator), RpoN (σ^54^), and the FlhF GTPase (Balaban et al., 2009; Joslin and Hendrixson, 2009, 2008). RpoN promotes transcription of class II genes including rod and hook components and the minor flagellin FlaB. In addition, RpoN activates transcription of FliA (σ^28^) and FlgM (Gilbreath et al., 2011) (**Fig. 1A**). Similar to Gammaproteobacteria, *H. pylori* and *C. jejuni* use a FlgM-FliA checkpoint inhibition strategy between expression of class II and class III genes to guarantee proper flagellar assembly (Josenhans et al., 2002; Wösten et al., 2010). Late flagellar genes include the major flagellin FlaA and other virulence-associated factors, such as flagellar co-expressed determinant (Fed) proteins (Barrero-Tobon and Hendrixson, 2012; Burnham and Hendrixson, 2018).

**Figure 1.**
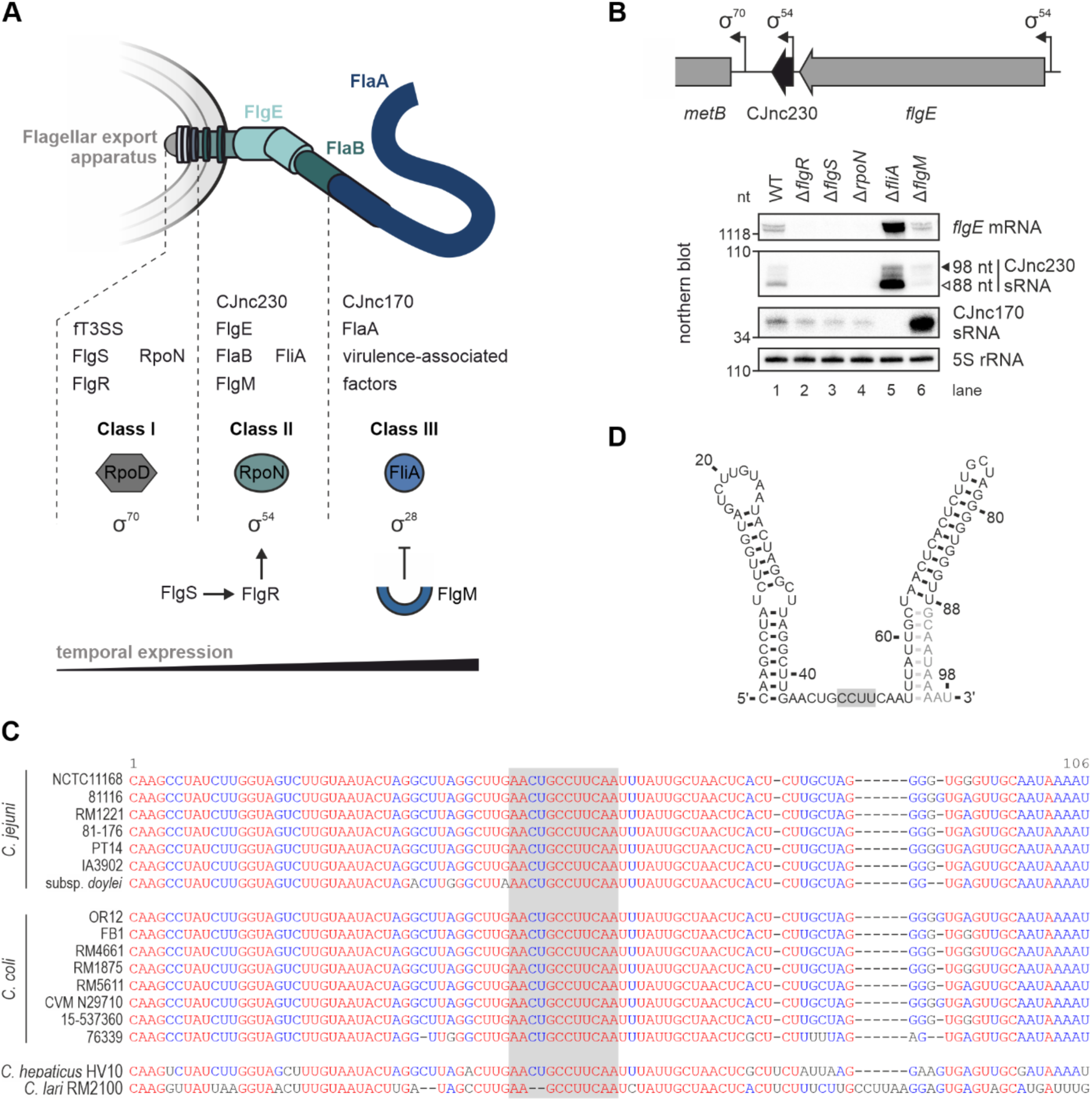
CJnc230 sRNA is transcribed in an RpoN-dependent manner and conserved in diverse *Campylobacter* species. **(A)** *C. jejuni* flagellum structure with flagellar export apparatus, hook (FlgE), filament components (FlaB & FlaA), and CJnc230/CJnc170 sRNAs. Below: temporal expression control by three sigma factors (RpoD, RpoN, FliA) of three classes of flagellar genes. Only proteins and sRNAs mentioned in the text are indicated. fT3SS: flagellar type three secretion system. **(B)** (*Upper*) Genomic location of CJnc230 (black arrow) in *C. jejuni* strain NCTC11168 downstream of the RpoN (σ^54^)-dependent *flgE* (Cj1729c) gene. Bent arrows: transcriptional start sites (TSSs) with associated promoter motifs (Dugar et al., 2013; Porcelli et al., 2013). (*Lower*) Northern blot analysis of total RNA from *C. jejuni* wildtype (WT) and flagella regulator deletion mutants at exponential growth phase. Black triangle: 98-nt previously annotated full-length sRNA (Dugar et al., 2013). White triangle: 3’-truncated abundant 88-nt version identified in this study. The *flgE* mRNA was detected with CSO-5136, CJnc230 sRNA was probed with CSO-0537, CJnc170 sRNA with CSO-0182, and 5S rRNA (CSO-0192) was detected as a loading control. **(C)** Alignment of corresponding 98-nt CJnc230 sequences from multiple *Campylobacter* species and strains using MultAlin (Corpet, 1988). Gray box: single-stranded RNA region (see panel **D**). **(D)** Secondary structure of full-length CJnc230 (98 nt) predicted by RNAfold (Gruber et al., 2008). Black residues: 3’-truncated 88-nt version. Gray box: putative anti-Shine-Dalgarno motif.

In contrast to the well studied hierarchical transcriptional control of flagellar biogenesis in diverse bacterial species, much less is known about the role of post-transcriptional regulation via small regulatory RNAs (sRNAs) within this cascade. These riboregulators mostly act via base-pairing interactions with target mRNAs and play important roles in bacterial adaptation to diverse environmental stresses and fine-tuning of multiple physiological processes, including virulence (Hör et al., 2020; Wagner and Romby, 2015; Westermann, 2018). So far, only a few gammaproteobacterial sRNAs are known to modulate flagellar biogenesis, mainly targeting the mRNA encoding FlhDC (De Lay and Gottesman, 2012; Mika and Hengge, 2013; Thomason et al., 2012) and even less is known about the role of sRNAs in the flagellar biosynthesis cascade outside of *E. coli* and *Salmonella*.

Several sRNA candidates identified based on global transcriptome analyses of *C. jejuni* have conserved motifs for flagellar sigma factors in their promoters (Dugar et al., 2013; Le et al., 2015; Porcelli et al., 2013), indicating they could be co-expressed and play a role in flagellar assembly. Here, we demonstrate that two distinct sRNAs are co-regulated with middle and late flagellar genes and post-transcriptionally fine-tune hierarchical control of flagellar gene expression in *C. jejuni*. CJnc230 sRNA is co-transcribed with the flagellar hook structural protein FlgE and processed by three ribonucleases (RNases). CJnc230 directly represses mRNAs encoding Cj1387c, a putative regulator of flagella-flagella interactions (CfiR) (Reuter et al., 2015), as well as the anti-sigma factor FlgM. Overexpression of CJnc230 increases flagellar length and motility via translational inhibition of *flgM*, thereby de-repressing FliA-dependent late flagellar genes, including the major flagellin FlaA. While CJnc230 transcription is dependent on RpoN, we show FlgM repression also induces the FliA-dependent sRNA CJnc170, which has an opposite effect on filament length and swimming. Based on these observations, we suggest renaming CJnc230 and CJnc170 to FlmE and FlmR (flagellar length and <u>m</u>otility <u>e</u>nhancer/<u>r</u>epressor), respectively. Due to their impact on the assembly of the flagellar complex, they affect a major virulence trait of this bacterium.

## Results

### RpoN is required for expression of the conserved *C. jejuni* sRNA CJnc230

While RNA-seq-based transcriptome analyses revealed diverse sRNA candidates in *C. jejuni* (Dugar et al., 2013; Porcelli et al., 2013; Taveirne et al., 2013), their functions and cellular targets remain largely unknown. To explore sRNAs that might regulate key *C. jejuni* phenotypes such as motility, we examined expression differences of several potential flagella co-regulated sRNAs in deletion mutants of characterized flagella regulatory genes by northern blotting. sRNAs were selected based on flagellar sigma factor promoter motifs or close vicinity to flagella structural genes. This confirmed a previously described dependency of CJnc170 sRNA on FliA (Le et al., 2015) and revealed that expression of CJnc230 sRNA, which is encoded downstream of the gene encoding the flagellar hook protein FlgE (Cj1729c) (Dugar et al., 2013; Porcelli et al., 2013), is strongly affected by deletion of several regulators involved in flagellar biosynthesis (**Fig. 1B**). CJnc230 expression was completely lost when the gene encoding RpoN or components of its upstream activating FlgRS two-component system were deleted and increased in a FliA mutant (**Fig. 1B**). Our previous comparative differential RNA-seq (dRNA-seq) identified CJnc230 as a conserved ∼98-nt long sRNA in *C. jejuni* strains NCTC11168, 81-176, 81116, and RM1221 (Dugar et al., 2013). While CJnc230 overlaps with the predicted open reading frame Cj1728c, we confirmed CJnc230 to be non-coding based on ribosome profiling (Froschauer et al., 2022). As CJnc230 expression was lost in the Δ*rpoN* mutant and transcription of the upstream encoded *flgE* is dependent on RpoN (Chaudhuri et al., 2011; Lertsethtakarn et al., 2011) (**Fig. 1B**), we hypothesized that CJnc230 might be a processed sRNA that is co-transcribed with *flgE.* In line with this, deletion of *fliA* strongly upregulated both *flgE* and two species (98/88 nt) of CJnc230 (**Fig. 1B**). This upregulation is consistent with previous reports for several other RpoN-dependent *C. jejuni* flagellar genes (Carrillo et al., 2004; Kamal et al., 2007), although the underlying mechanism is so far unclear. Moreover, in contrast to the strong upregulation of FliA-dependent CJnc170, CJnc230 and *flgE* levels slightly decreased in a mutant lacking the anti-σ^28^ factor FlgM, further supporting their co-expression.

To further examine CJnc230 conservation, we performed sequence conservation analyses using blastn and GLASSgo (Lott et al., 2018). This revealed CJnc230 is highly conserved in diverse *C. jejuni* and *C. coli* strains (**Fig. 1C**), the two most common agents of *Campylobacter*-mediated gastroenteritis in humans (Janssen et al., 2008). Manual inspection revealed highly similar sequences also in more distant *C. hepaticus* (85.9% identity) and *C. lari* (69.6% identity) compared to *C. jejuni* strain NCTC11168. Secondary structure predictions for the full-length 98-nt sRNA with RNAfold (Gruber et al., 2008) revealed two stem-loops flanking a single-stranded stretch of 12 nt (**Fig. 1D**). This single-stranded region contains a conserved sequence with a potential anti-Shine-Dalgarno sequence (CCUU), complementary to the *C. jejuni* ribosome binding site (RBS) consensus motif (Froschauer et al., 2022), suggesting a role as a *trans*-acting mRNA regulator. In all species that a homologous CJnc230 sequence was identified in, the sRNA was encoded downstream of the corresponding *flgE* ortholog (**Figs. 1B** **& S1A**), which, together with its dependence on RpoN, points to a possible role of CJnc230 in *Campylobacter* flagellar biogenesis.

### RNase III, RNase Y, and PNPase are involved in processing of CJnc230

CJnc230 seemed to be coupled with *flgE* mRNA levels and was detected as multiple species (∼98 and ∼88 nt) on northern blots (**Fig. 1B**). Thus, we investigated its possible co-transcription with *flgE* and subsequent processing by RNases. Unlike other Gram-negative bacteria, *C. jejuni* does not encode RNase E, responsible for processing many 3’ untranslated region (UTR)-derived sRNAs in Gammaproteobacteria (Miyakoshi et al., 2015; Ponath et al., 2022), and instead encodes RNase Y (Parkhill et al., 2000). Double-strand-specific RNase III is also emerging as a key player in sRNA processing and regulation (Altuvia et al., 2018; Lioliou et al., 2012; Mediati et al., 2022), and important for ribosomal and CRISPR RNA biogenesis in *C. jejuni* (Dugar et al., 2018, 2013; Haddad et al., 2013). We measured CJnc230 expression in total RNA from a panel of RNase-deficient *C. jejuni* deletion mutant strains by northern blot (**Fig. S1B**), which revealed that RNase III (*rnc*, Cj1635c), RNase Y (*rny*, Cj1209), and the 3’-5’ exoribonuclease polynucleotide phosphorylase (PNPase, *pnp*, Cj1253) affect its expression (**Figs. 2A** **& S1B**). In Δ*rnc*, mature CJnc230 was not observed, and instead a transcript longer than 1,000 nt, potentially comprising *flgE*-CJnc230, was detected (**Fig. 2A**, lane 2). Upon deletion of RNase Y, several transcripts of approximately 200-300 nt appeared in addition to the mature sRNA bands present in wildtype (WT) (**Fig. 2A**, lane 3), suggesting RNase Y also affects CJnc230 processing. Accumulation of the most abundant WT RNA species (approximately ∼88 nt) was increased in the PNPase deletion mutant compared to WT (**Fig. 2A**, lane 4, white triangle). In an *rny*/*pnp* double deletion, additional species between 110 and 300 nt were also detected (**Fig. 2A**, lane 5). An oligonucleotide binding downstream of the annotated sRNA 3’ end (Dugar et al., 2013) (3’-extended, CSO-4297) revealed that RNase Y is involved in 3’-end maturation of CJnc230 or degradation of longer RNA species, as only the higher molecular weight fragments were detected (**Fig. 2A**, lane 3). PNPase seems to also contribute to processing of some 3’-extended transcripts, since the additional fragments between 200 and 300 nt were also observed in the *rny*/*pnp* double mutant when using CSO-4297 for hybridization (**Fig. 2A**, lane 5).

**Figure 2.**
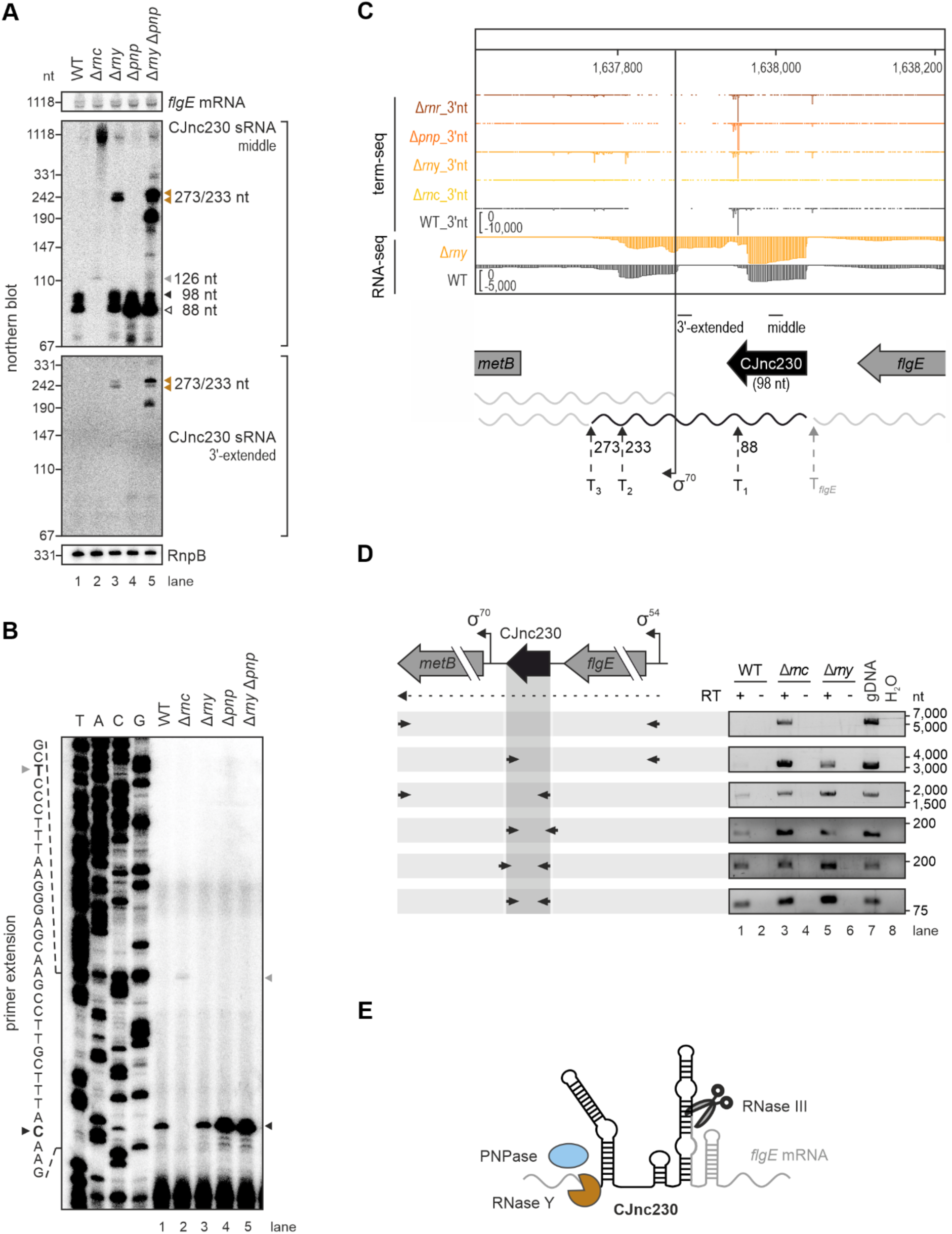
CJnc230 is co-transcribed with *flgE* and processed by RNase III, RNase Y, and PNPase. **(A)** Northern blot analyses of total RNA from *C. jejuni* WT and ribonuclease (RNase) deletion mutant strains in exponential phase. Colored triangles: prominent CJnc230 transcripts determined by termination site sequencing (term-seq) and/or primer extension. The *flgE* mRNA was detected with CSO-5136 (binding the coding sequence (CDS)), CJnc230 sRNA was probed with CSO-0537 (middle of sRNA) or CSO-4297 (3’-extended versions, binding ∼70 nt downstream of annotated 3’ end (Dugar et al., 2013)). RnpB RNA (CSO-0497) served as a loading control. Oligonucleotide binding positions are indicated in (**C**). **(B)** Primer extension analysis of CJnc230 5’ ends from *C. jejuni* WT and RNase deletion mutant strains in exponential phase using the CJnc230 probe used for northern blots (CSO-0537) binding in the middle of the sRNA. A sequencing ladder generated with this probe is partially indicated on the left. Black triangle: CJnc230 5’ end in the WT (C residue in bold on the left). Gray triangle: alternative 5’ end 28 nt upstream in Δ*rnc* (T residue in bold). **(C)** Term-seq and total RNA-seq cDNA coverages at the CJnc230 locus in *C. jejuni* WT and nuclease deletion mutants grown to exponential phase. One representative replicate each is shown. For term-seq libraries, coverage for the last base of each read was plotted (3’nt), while total RNA-seq tracks depict full read coverage. Bent arrow: *metB* transcriptional start site (TSS) from (Dugar et al., 2013). Dashed arrows: prominent CJnc230 3’ ends (T_1_, T_2_, and T_3_) with resulting sRNA lengths in nucleotides and RNA 3’ end downstream of *flgE* (T*_flgE_*). Oligonucleotide binding positions used for northern blot in (**A**) are indicated. **(D)** Reverse transcription-polymerase chain reaction (RT-PCR) analysis of total RNA from *C. jejuni* WT, Δ*rnc*, and Δ*rny* harvested at exponential growth phase. (*Left*) The *flgE*-CJnc230-*metB* locus. Arrows below: RT-PCR primers. Bent arrows: TSSs & promoter motifs (Dugar et al., 2013; Porcelli et al., 2013). (*Right*) Agarose gels of RT-PCR reactions performed in the presence (+) or absence (-) of reverse transcriptase (RT). Reactions with genomic DNA (gDNA) of WT or water (H_2_O) served as positive and negative controls, respectively. **(E)** Model of CJnc230 processing and maturation by three ribonucleases. RNase III potentially cleaves a predicted stem-loop in the *flgE*-CJnc230 transcript (**Fig. S3**), while RNase Y and PNPase are involved in maturation of the sRNA 3’ end (full-length CJnc230, 98 nt, colored in black) or in degradation of precursor transcripts.

To further investigate processing of CJnc230, we used primer extension to locate 5’ ends in the RNase mutant strains. This confirmed the previously annotated CJnc230 5’ end in the WT strain (**Fig. 2B**, lane 1, black triangle). This also showed that this end is potentially RNase III-dependent, as the 5’ end was shifted upstream by 28 nt in the *rnc* deletion mutant (**Fig. 2B**, lane 1 vs. 2, gray triangle), despite it being annotated as a transcriptional start site (TSS) (Dugar et al., 2013; Porcelli et al., 2013). A longer version of the sRNA (∼126 nt) was also detected in Δ*rnc* total RNA by northern blot (**Fig. 2A**, lane 2, gray triangle). So far, it was unclear whether this 28 nt-extended 5’ end originates from an independent promoter or from cleavage of the *flgE* promoter-derived transcript by a yet unknown RNase. All other nuclease deletion mutants tested did not show 5’ end differences from WT (**Fig. 2B**, lane 1 vs. lanes 3-5, black triangle), although higher levels of the mature CJnc230 5’ end were detected in strains without PNPase (**Fig. 2B**, lanes 4 and 5), comparable to northern blot observations (**Fig. 2A**). These results suggest that CJnc230 biogenesis involves three RNases: RNase III cleaves CJnc230 at the 5’ end, while RNase Y and PNPase affect 3’-end processing of the sRNA.

### Term-seq reveals the most abundant 3’ end of CJnc230

The CJnc230 expression pattern in WT bacteria on northern blots showed one highly abundant RNA species of lower molecular weight (∼88 nt) and several slightly longer transcripts (∼88-98 nt) (**Figs. 1B** **& 2A**, lane 1). Since primer extension analysis identified only one 5’ end in the WT background (**Fig. 2B**), this pattern likely arises from differences at the sRNA 3’ end. To follow this up, we globally identified RNA 3’ ends using termination site sequencing (term-seq) (Dar et al., 2016) in *C. jejuni* NCTC11168 WT, Δ*rnc*, Δ*rny*, Δ*pnp*, and Δ*rnr* strains. A mutant of RNase R (*rnr*, Cj0631c) was included, as this enzyme shows 3’-5’ exoribonuclease activity in *C. jejuni* (Haddad et al., 2014). Visual inspection of term-seq data in WT, Δ*rny*, Δ*pnp*, and Δ*rnr* revealed multiple CJnc230 3’ ends, resulting in transcripts from 87 to 98 nt in length according to the mature sRNA 5’ end (**Fig. 2C**). This 3’ end variability might explain the diversity of CJnc230 RNA species detected on northern blots (**Figs. 1B** **& 2A**, lane 1). The most prominent peak in WT, Δ*rny*, Δ*pnp*, and Δ*rnr* was located 88 nt downstream of the CJnc230 5’ end, consistent with the most abundant northern blot species and the primer extension-located 5’ end (**Figs. 1B** **& 2A, 2B**), and giving rise to a version of the sRNA with a predicted longer single-stranded stretch (**Fig. S2A**). In contrast to the increase of the 88-nt version, the abundance of longer CJnc230 species was reduced ∼2-fold over growth in WT (**Fig. S2B**, panel 3’ end oligo, lanes 1, 5, and 9). The variation in CJnc230 3’ ends and the sRNA 5’ end was also confirmed by circular Rapid Amplification of cDNA Ends (cRACE) (**Fig. S2C**). In agreement with northern blots, only low CJnc230 expression was observed in term-seq of Δ*rnc* bacteria and all prominent peaks were missing (**Fig. 2C**). Another peak was detected nine nucleotides upstream of the sRNA 5’ end in WT, which might represent a *flgE* transcript 3’ end. As this position was not immediately adjacent to the sRNA 5’ end, but was RNase III-dependent, this suggests that there is either further degradation of the *flgE* 3’ end or sRNA 5’ end.

Term-seq of Δ*rny* revealed two additional peaks 233 and 273 nt downstream of the mature sRNA 5’ end that was determined by primer extension (**Fig. 2C**). These positions (T2 and T3), also reflected by additional total RNA-seq coverage downstream of the sRNA 3’ end in the Δ*rny* library compared to WT (**Fig. 2C**), were consistent with two additional CJnc230 bands detected in this mutant on northern blots, thus confirming these as 3’-extended transcripts (**Fig. 2A**, lanes 3 & 5, orange triangles).

### CJnc230 is co-transcribed with *flgE* and *metB* mRNAs

The similar RpoN dependence of *flgE* mRNA and CJnc230 on northern blot (**Fig. 1B**) and the RNase III cleavage at the CJnc230 5’ end hinted towards their co-transcription. We next used reverse transcription-PCR (RT-PCR) with different primer combinations to determine if they are part of the same RNA. Indeed, we found that CJnc230 is transcribed together with *flgE,* and also with the downstream gene *metB* (O-acetylhomoserine (thiol)-lyase, Cj1727c), which also has its own dedicated RpoD-dependent TSS (**Fig. 2D**). However, *flgE*-CJnc230-*metB* and *flgE*-CJnc230 transcripts were hardly detectable in the WT strain when RNase III was present (**Fig. 2D**, two upper boxes). RNase Y appears to play a role in 3’-maturation of CJnc230 (**Fig. 2A**), but it also seems to destabilize the *flgE*-CJnc230 transcript, as this fragment could be detected in the Δ*rny* background where RNase III is present (**Fig. 2D**, second box from top, lanes 1 and 5). Oligonucleotides binding the CJnc230 5’ or 3’ end were used to detect the sRNA only, as a control (**Fig. 2D**, last box). Interestingly, a 5’-extended sRNA was also detected in the WT background, arguing for a version originating upstream of the RNase III cleavage site (**Fig. 2D**, third last box, lane 1). RNA species, which are extended further downstream beyond the CJnc230 3’ end, were detectable in all strain backgrounds, either only ∼70 nt downstream of the 3’ end of the full-length sRNA (**Fig. 2D**, second last box) or until the *metB* stop codon (**Fig. 2D**, third box from top). A long stem-loop structure was predicted using the *flgE*-CJnc230 transcript as input (**Fig. S3**), potentially serving as template for RNase III cleavage between positions 2,732 and 2,733 (between A-U and C-G base pairs) relative to the *flgE* TSS. Or CJnc230 maturation might also include other so-far unknown RNA partners binding to CJnc230, thereby promoting RNase III cleavage at the sRNA 5’ end.

In summary, based on northern blotting, primer extension, term-seq, cRACE, and RT-PCR, we could show that CJnc230 is generated by a complex biogenesis. This involves transcription together with the RpoN-dependent gene *flgE*, cleavage downstream of the *flgE* coding sequence (CDS) by RNase III, and 3’ end maturation/degradation by RNase Y and PNPase (**Fig. 2E**).

### CJnc230 is also transcribed from an RpoD-promoter downstream of the *flgE* CDS

To determine whether the alternative CJnc230 5’ end observed in Δ*rnc* (**Fig. 2B**) originated from an independently transcribed version of the sRNA, in addition to processing from the *flgE* promoter-derived transcript, we globally mapped primary transcript 5’ ends with dRNA-seq in NCTC11168 WT and Δ*rnc*. This revealed a potential alternative TSS in Δ*rnc* with a putative RpoD-dependent promoter motif (5’-TATTGT-3’; -41 to -36 relative to annotated 5’ end (Dugar et al., 2013)) (**Fig. S4**), upstream of the annotated CJnc230 5’ end that was confirmed by primer extension in WT bacteria (**Fig. 2B**). This alternative TSS was consistent with the 28 nt-extended 5’ end detected in Δ*rnc* by primer extension (**Fig. 2B**). In the dRNA-seq experiment performed here we did not observe enrichment of the annotated CJnc230 5’ end (Dugar et al., 2013) in the TEX-treated WT sample compared to the untreated sample. This suggests processing of the sRNA 5’ end, even though we also found a conserved putative RpoN-dependent motif upstream of it (5’-GG TTGCTT-3’; position -18 to -3 relative to annotated 5’ end) (**Fig. S4**).

To determine if there is an active promoter in this region, we generated a transcriptional reporter by fusing 150 nt upstream of the annotated sRNA 5’ end to the unrelated *metK* RBS and a superfolder GFP (sfGFP) gene (Pédelacq et al., 2006) (**Fig. S5A**). Western blot analysis of whole cell lysates confirmed that this region alone could drive transcription of the reporter. This transcription was independent of flagellar sigma factors (RpoN or FliA) (**Fig. S5A**, upper panel of western blot, lanes 1 to 4), unlike the *flgE* promoter, which we confirmed is RpoN dependent (lower panel of western blot). This suggested that CJnc230 might also be transcribed from its own RpoD-dependent promoter in addition to transcription together with upstream *flgE*. Interestingly, we also observed that deletion of *fliA* increased activity of P*flgE*, in line with increased *flgE* mRNA abundance in Δ*fliA* (**Fig. 1B**). This suggests increased *flgE* levels might originate from transcriptional feedback on class II flagellar genes via a yet unknown mechanism, as observed previously in microarray analyses (Carrillo et al., 2004; Kamal et al., 2007).

Primer extension analysis located a 5’ end in Δ*rnc* that is in line with the alternative CJnc230 TSS 28 nt upstream of the 5’ end in WT (**Figs. 2B** **& S5B**, lane 2, gray triangle), supporting the presence of an independently transcribed sRNA in addition to the *flgE* promoter-derived version. Analysis of strains deleted for *rpoN*, *flgE* (including its promoter), or both, also confirmed that the sRNA can be transcribed independently of *flgE*, possibly from its own RpoD-dependent promoter (black triangle). However, this transcript also appears to be further processed, as the 5’ end is the same as in WT bacteria (**Fig. S5B**, lanes 6 to 8, black triangle). Strikingly, the RpoD-dependent version of CJnc230 seemed to be independent of RNase III, as the WT 5’ end and sRNA expression was detected in a *flgE/rnc* double mutant by primer extension or northern blot, respectively (**Fig. S5B, C**; lane 9). Northern blot analysis confirmed the 28-nt 5’-extended transcript in Δ*rnc* samples (**Figs. 2B** **& S5C**, lane 2, gray triangle), as well as very low levels of sRNA expression in *rpoN* and *flgE* single or double deletion mutants with a similar 88-nt length as in WT bacteria (**Fig. S5C**, lanes 6 to 8, white triangle). Taken together, our data suggest that CJnc230 can also be generated by an additional biogenesis pathway, which is independent of *flgE* transcription and relies on its own RpoD-dependent promoter. However, this generates very low levels of CJnc230 under the conditions examined. The 126 nt-long precursor transcript generated from the promoter also seems further processed by a yet unknown RNase, as the sRNA 5’ end originating from either biogenesis pathway is the same.

### CJnc230 directly represses translation of Cj1387c and *flgM* mRNAs

Next, to investigate CJnc230 function and cellular targets in *C. jejuni*, we constructed deletion (Δ), complementation (C), and overexpression (OE) mutants in strain NCTC11168. Approximately 14- to ∼26-fold overexpression of CJnc230 was achieved during different growth phases (**Fig. S2B**) by fusing the full-length sRNA (98 nt) to the RpoD-dependent *porA* promoter at the unrelated *rdxA* locus (Ribardo et al., 2010). No significant difference in growth was observed for any of the mutant strains compared to WT bacteria (**Fig. S6A**). However, inspection of whole cell protein samples of the mutant strains by SDS-PAGE revealed slight upregulation of bands consistent with the sizes of FlaA and FlaB flagellins (Radomska et al., 2017) upon overexpression of CJnc230 (**Fig. S6B**), suggesting CJnc230 might play a role in regulation of *C. jejuni* flagellar biogenesis.

To identify potential direct targets of CJnc230 in *C. jejuni* strain NCTC11168, we performed *in-silico* target predictions with IntaRNA (Mann et al., 2017) for both the 98-nt full-length and the 88-nt version of the sRNA. Three quarters of the predicted targets for full-length CJnc230 overlapped with predictions for the shorter, more abundant 88-nt version, which has an almost two-fold longer single-stranded RNA region and also larger pool of predicted targets (**Tables S1** & **S2,** **Figs. 1D** **& S2A**). The top 10 overlapping candidates included mRNAs encoding flagella-related proteins, of which *flgM* and Cj1387c mRNAs were predicted to interact with the single-stranded region of CJnc230 at their RBS (**Fig. 3A**), consistent with repression via competition for ribosome binding. While the anti-FliA factor FlgM (Cj1464) regulates late flagellar gene expression by sequestering σ^28^ (FliA) (Wösten et al., 2010) (**Fig. 1A**), Cj1387c was previously suggested to play a role in flagella-flagella interactions and thus named *Campylobacter* flagella interaction regulator (CfiR) (Reuter et al., 2015). In-line probing (Regulski and Breaker, 2008) supported the predicted structure of CJnc230 (**Fig. 1D**) and confirmed its interaction with the two targets (**Fig. 3B**). Addition of increasing concentrations of *in-vitro* transcribed, unlabeled mRNA fragments including their RBS protected the single-stranded region of 5’-radiolabeled CJnc230 from spontaneous degradation (**Fig. 3B**, lanes 4 to 7 for Cj1387c and 8 to 11 for *flgM*). In contrast, a mutant sRNA with two base exchanges in the predicted interaction region (M1) was not protected, even with high concentrations of Cj1387c and *flgM* transcripts (**Fig. 3B**, lanes 15 to 17). These results were validated by a reciprocal experiment with 5’-radiolabeled Cj1387c and *flgM* mRNA leaders and unlabeled CJnc230 (**Fig. S7A**).

**Figure 3.**
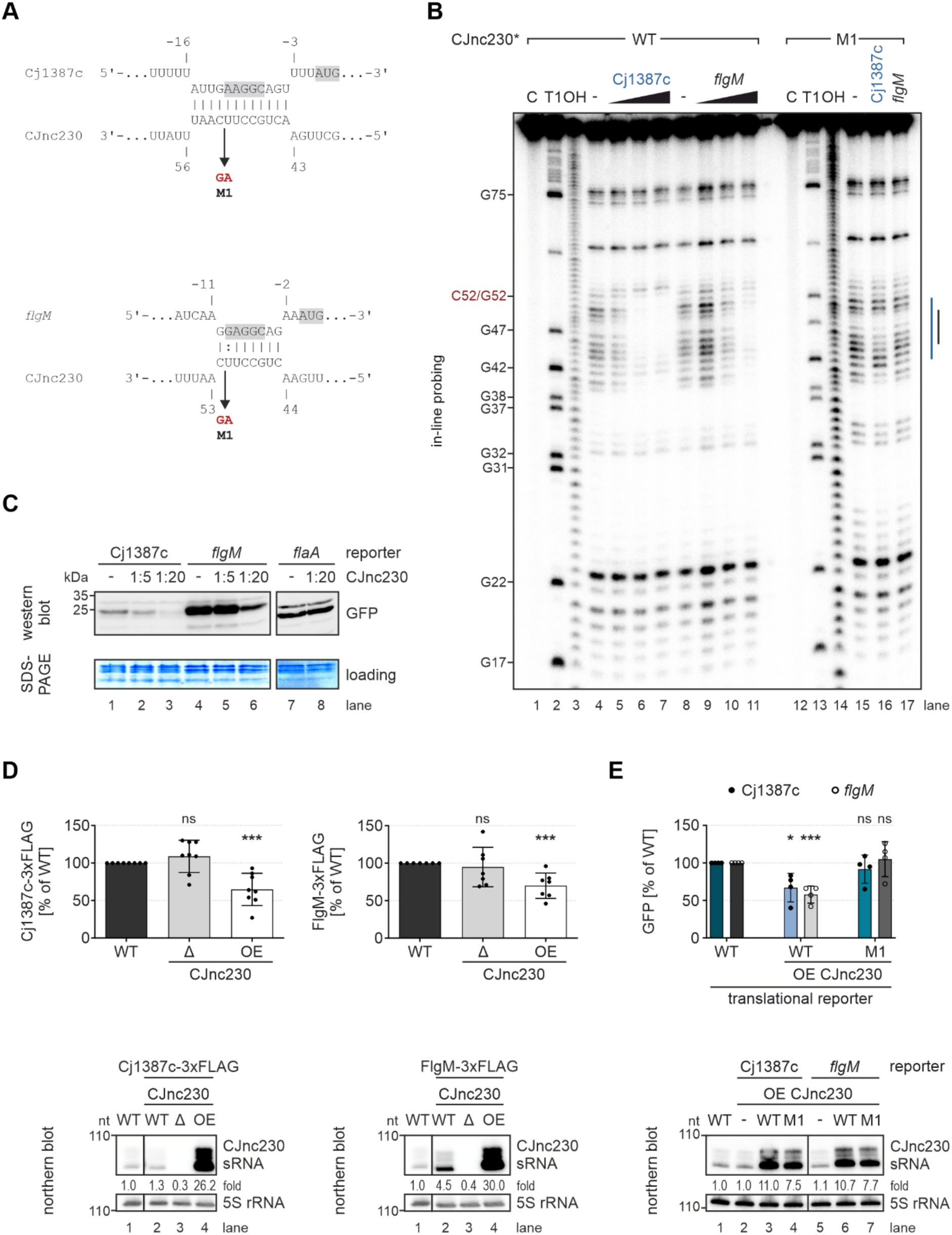
CJnc230 interacts with the ribosome binding site (RBS) of Cj1387c and *flgM* mRNAs to repress their translation. **(A)** Predicted interactions between the single-stranded region of CJnc230 sRNA and Cj1387c or *flgM* mRNAs (IntaRNA) (Mann et al., 2017). Gray boxes: RBS and start codons. Arrow/M1: sRNA mutations. **(B)** In-line probing of 0.2 pmol ^32^P-5’-end-labeled (marked with *) WT CJnc230 sRNA in the absence or presence of 0.02/0.2/2 pmol unlabeled Cj1387c or *flgM* WT leaders. Interaction sites for Cj1387c (blue) and *flgM* (black) are indicated on the right. 5’-end-labeled mutated sRNA (M1; C52G mutation in red on the left; 0.2 pmol) was incubated with 2 pmol of WT mRNA leaders. C - untreated control; T1 ladder - G residues (indicated on the left); OH - all positions (alkaline hydrolysis). **(C)** *In-vitro* translation of *in-vitro* transcribed mRNAs of Cj1387c/*flgM*-sfGFP reporters (4 pmol, 5’UTR and first 10 codons fused to *sfgfp*) in an *E. coli* cell-free system -/+ CJnc230 (1:5 or 1:20) detected by western blot. *flaA* 5’UTR reporter: negative control, separate western blot. PageBlue staining of the gel after transfer served as a loading control. **(D)** (*Upper*) Protein levels of C-terminally FLAG-tagged Cj1387c (*left*) and FlgM (*right*) in exponentially grown *C. jejuni* measured by western blot. Mean of independent replicates (n = 8 for Cj1387c-3xFLAG; n = 7 for FlgM-3xFLAG), error bars depict the standard deviation. ***: *p* < 0.001, ns: not significant, two-tailed Student’s *t*-test was used to compare the respective mutant to WT. (*Lower*) Northern blot analysis of total RNA from *C. jejuni* WT and CJnc230 deletion (Δ) and overexpression (OE) mutants in epitope-tagged backgrounds grown to exponential phase. CJnc230 was detected with CSO-0537 and 5S rRNA (CSO-0192) served as a loading control. Fold changes of CJnc230 expression relative to WT and normalized to 5S rRNA are indicated. Images in (**D**) were cut between lanes 1 and 2. **(E)** (*Upper*) Western blot analysis of *C. jejuni* translational reporters of Cj1387c (filled circles) and *flgM* (open circles) 5’UTRs (and first 10 codons) fused to *sfgfp*, harvested at exponential growth phase. sRNA WT or M1 mutant OE strains are compared to the 5’UTR reporters without sRNA overexpression. Mean of independent replicates (n = 4), error bars depict the standard deviation. ***: *p* < 0.001, *: *p* < 0.05, ns: not significant, two-tailed Student’s *t*-test. (*Lower*) Northern blot analysis of total RNA from *C. jejuni* WT and CJnc230 (WT/M1) OE mutants in translational reporter backgrounds grown to exponential phase. Expression of CJnc230 sRNA was detected with CSO-0537 and 5S rRNA (CSO-0192) served as a loading control. Fold changes of CJnc230 expression relative to the WT and normalized to 5S rRNA are indicated. Images in (**E**) were cut between lanes 4 and 5.

Next, we confirmed direct translational repression of *flgM* and Cj1387c mRNAs by CJnc230 in an *in-vitro* system. Translational reporters for each target were generated by fusing their 5’UTR and first ten codons to *sfgfp* and *in-vitro* transcribed using T7 RNA polymerase. We observed dose-dependent translational repression of both reporters by CJnc230 on western blots (**Fig. 3C**, lanes 1 to 6). As a negative control, no repression was observed for a *flaA* translational reporter, which is not predicted to be a target of CJnc230 (**Fig. 3C**, lanes 7 and 8; **Tables S1 & S2**). When sRNA excess was further increased to 50-fold, a complete inhibition of Cj1387c and *flgM* reporter translation was observed (**Fig. S7B**).

To demonstrate that CJnc230 can repress translation of Cj1387c and *flgM in vivo*, we chromosomally tagged both genes with a 3xFLAG epitope at their C-terminus. Western blot analysis of whole cell lysates revealed that while deletion of CJnc230 did not significantly affect protein levels, ∼26-30-fold overexpression of CJnc230 led to a significant reduction of Cj1387c- and FlgM-3xFLAG by ∼35% and ∼30%, respectively (**Fig. 3D**). Additionally, translational *sfgfp* reporters driven from an unrelated promoter (*metK*), fused to the 5’UTRs and first 10 codons of both targets were also downregulated in *C. jejuni* upon overexpression of the sRNA (**Fig. 3E**). GFP expression was significantly reduced only when WT CJnc230 was overexpressed (by ∼42% for *flgM* and ∼33% for Cj1387c), but not when overexpressing the M1 mutant, although both sRNA versions were overexpressed to similar levels (**Fig. 3E**). In summary, data from both *in-vitro* and *in-vivo* approaches validated post-transcriptional repression of Cj1387c and *flgM* by CJnc230, mediated via direct binding to the mRNA RBSs.

### CJnc230-mediated FlgM repression leads to increased FlaA expression, filament length, and motility

Next, we investigated the effect of CJnc230-mediated FlgM repression on flagellar genes downstream of FlgM. SDS-PAGE analysis of total proteins suggested that CJnc230 overexpression upregulates either the FlaA (FliA-dependent) or FlaB (RpoN-dependent) flagellins, which have a similar size (**Fig. S6B**). To identify if FlaA or FlaB are affected by CJnc230, we measured levels of 3xFLAG-tagged flagellins in CJnc230 mutants by western blot analysis. While deletion of CJnc230 had no significant effect on FlaA- or FlaB-3xFLAG levels, ∼32-fold overexpression of the sRNA significantly upregulated epitope-tagged FlaA levels by ∼118% compared to the WT background (**Figs. 4A** **& S8A**). As FlaB-3xFLAG levels remained unchanged upon CJnc230 overexpression (**Fig. S8A**), this argues for a specific effect of CJnc230 on class III flagellar genes via repression of FlgM. As a control, we tested if expression of the alternative sigma factors was altered by CJnc230 inhibition of *flgM*. Western blot analysis of CJnc230 deletion or overexpression strains in RpoN-3xFLAG or FliA-3xFLAG backgrounds showed that tagged protein levels were not affected by the sRNA (**Fig. S8B**).

**Figure 4.**
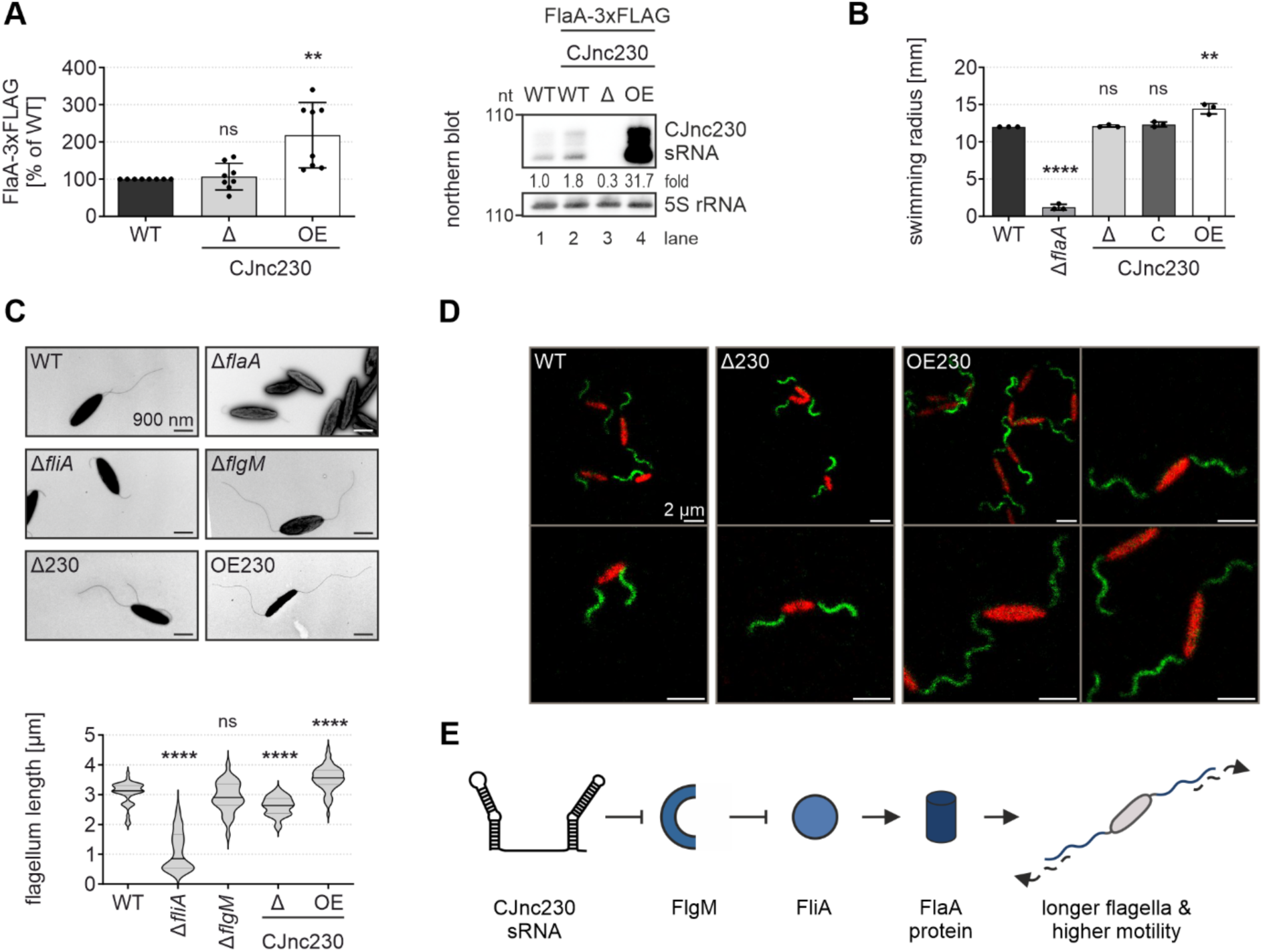
CJnc230 overexpression increases major flagellin levels, filament length, and motility. **(A)** (*Left*) FlaA-3xFLAG levels in *C. jejuni* WT, CJnc230 deletion (Δ), and overexpression (OE) mutants harvested at exponential growth phase and quantified by western blotting. Mean of independent replicates (n = 8), error bars depict the standard deviation. **: *p* < 0.01, ns: not significant, two-tailed Student’s *t*-test was used to compare the respective mutant to WT. (*Right*) Northern blot analysis of total RNA from *C. jejuni* WT and CJnc230 mutants in the FlaA-3xFLAG background grown to exponential phase. CJnc230 sRNA was detected with CSO-0537 and 5S rRNA (CSO-0192) served as a loading control. Fold changes of CJnc230 expression relative to WT and normalized to 5S rRNA are indicated. **(B)** Motility assays of WT and mutant strains with ΔCJnc230, complementation in *trans* (C CJnc230), and OE CJnc230 in 0.4% soft agar BB plates. Δ*flaA*: non-motile control. Error bars: standard deviation of the mean of independent replicates (n = 3). ****: *p* < 0.0001, **: *p* < 0.01, ns: not significant, two-tailed Student’s *t*-test vs. WT. A representative plate image is shown in **Figure S10A**. **(C)** Transmission electron microscopy (TEM) of *C. jejuni* WT, Δ*flaA*, Δ*fliA*, Δ*flgM*, and CJnc230 deletion (Δ230) or overexpression (OE230) mutants. (*Upper*) Representative TEM images at 6,000 x magnification, scale bar = 900 nm. (*Lower*) Violin plots depict the median (black) and quartiles (gray) of at least 30 flagellum length measurements per strain. ****: *p* < 0.0001, ns: not significant, two-tailed Student’s *t*-test was used to compare the respective mutant to WT. **(D)** Representative confocal microscopy images of *C. jejuni* expressing a second copy of *flaA* with a S395C mutation under control of its native promoter introduced at the *rdxA* locus. Bacteria were harvested at exponential growth phase and flagella were stained with DyLight™ 488 maleimide (green), while cell bodies were counterstained with eFluor™ 670 (red). Scale bar = 2 µm. **(E)** Model of CJnc230 function in *C. jejuni* strain NCTC11168. The sRNA represses translation of *flgM*, thereby de-repressing FliA-dependent genes, *e.g.*, *flaA*. This leads to increased FlaA protein levels, filament length, and motility.

Because the major flagellin FlaA is the main component of the *Campylobacter* flagellar filament and is required for motility (Burnham and Hendrixson, 2018), we compared the swimming behavior of CJnc230 mutants to WT. While deletion of CJnc230 did not affect motility, overexpression of CJnc230 slightly, but significantly, increased mean swim halo radii on soft agar compared to the WT strain (12 mm for WT vs. 14.4 mm for OE CJnc230) (**Figs. 4B** **& S10A**). Additionally, as *flgM* mutants in different bacterial species, including another *C. jejuni* strain grown under different conditions, express longer flagella or have twice the number of flagella per cell (Correa et al., 2004; Kutsukake and Iino, 1994; Wösten et al., 2010), we determined whether flagellar morphology was also affected by CJnc230. Transmission electron microscopy revealed significantly shorter flagella in the CJnc230 deletion background and significantly longer flagella in the sRNA overexpression mutant compared to the mean filament length in WT bacteria (WT: 3,064 nm; ΔCJnc230: 2,614 nm; OE CJnc230: 3,549 nm) (**Fig. 4C** **& Table S3**). Increased flagellum length upon overexpression of CJnc230 was further validated by using a mutated version of FlaA (S395C) in combination with a maleimide-conjugated fluorophore, allowing for staining of the flagellar filament. Employing confocal microscopy, we observed that a larger portion of OE CJnc230 mutant flagella had two, rather than one and a half, turns compared to either ΔCJnc230 or WT backgrounds (**Fig. 4D**). Bacteria with a *fliA* deletion were either aflagellate or had shorter flagella, possibly consisting of FlaB minor flagellin units only (class II, RpoN-dependent). This resulted in a mean flagellum length of 1,100 nm (**Fig. 4C** **& Table S3**), which is similar to previous reports for this mutant (Dugar et al., 2016). We did not observe a significantly different mean flagellum length of a Δ*flgM* strain compared to WT (2,926 nm) (**Fig. 4C** **& Table S3**). This indicates that there might be additional mechanisms compensating for the complete loss of FlgM in the deletion strain that are absent in the CJnc230 overexpression strain, or that additional CJnc230 targets affect flagellum length. Taken together, our data suggest that overexpression of CJnc230 increases flagella length via repression of *flgM* and upregulation of FlaA (**Fig. 4E**).

### CJnc230 overexpression increases transcription of class III genes, including the sRNA CJnc170

Since we observed that CJnc230 overexpression increased *C. jejuni* NCTC11168 motility and flagellar length (**Fig. 4B-D**), we hypothesized that these phenotypes are due to increased FlaA levels as a result of FlgM repression (and FliA de-repression). Thus, to test whether CJnc230 overexpression de-represses transcription of FliA-dependent genes, we measured *flaA* mRNA levels by northern blot in WT and CJnc230 mutants. In addition, we also examined levels of CJnc170 sRNA, which was previously shown to be FliA-dependent and suggested to control multiple RpoN-dependent genes, such as *flgE* (Cj1729c), *flaB* (Cj1338c), and *flgP* (Cj1026c) (Le et al., 2015). Compared to WT, ΔCJnc230 did not show different levels of either *flaA* mRNA or CJnc170 sRNA (**Fig. 5A**, compare lanes 1 and 2). However, overexpression of CJnc230 increased the abundance of both transcripts (**Fig. 5A**, compare lanes 1 and 4). Transcript levels of the RpoN-dependent class II gene *flgE* were slightly decreased in the CJnc230 overexpression mutant and remained unchanged in deletion and complementation strains. Although expression of *flgE* mRNA (and CJnc230) was lost in the *rpoN* deletion mutant, confirming this transcript as class II gene, it was increased in the Δ*fliA* strain (**Fig. 5A**, compare lanes 1 and 6), suggesting that there is transcriptional feedback on class II flagellar genes (**Fig. S5A**, lower panel). In contrast, *flaA* mRNA and CJnc170 sRNA were downregulated in Δ*rpoN* but completely lost upon deletion of *fliA*, as expected for class III genes (**Fig. 5A**).

**Figure 5.**
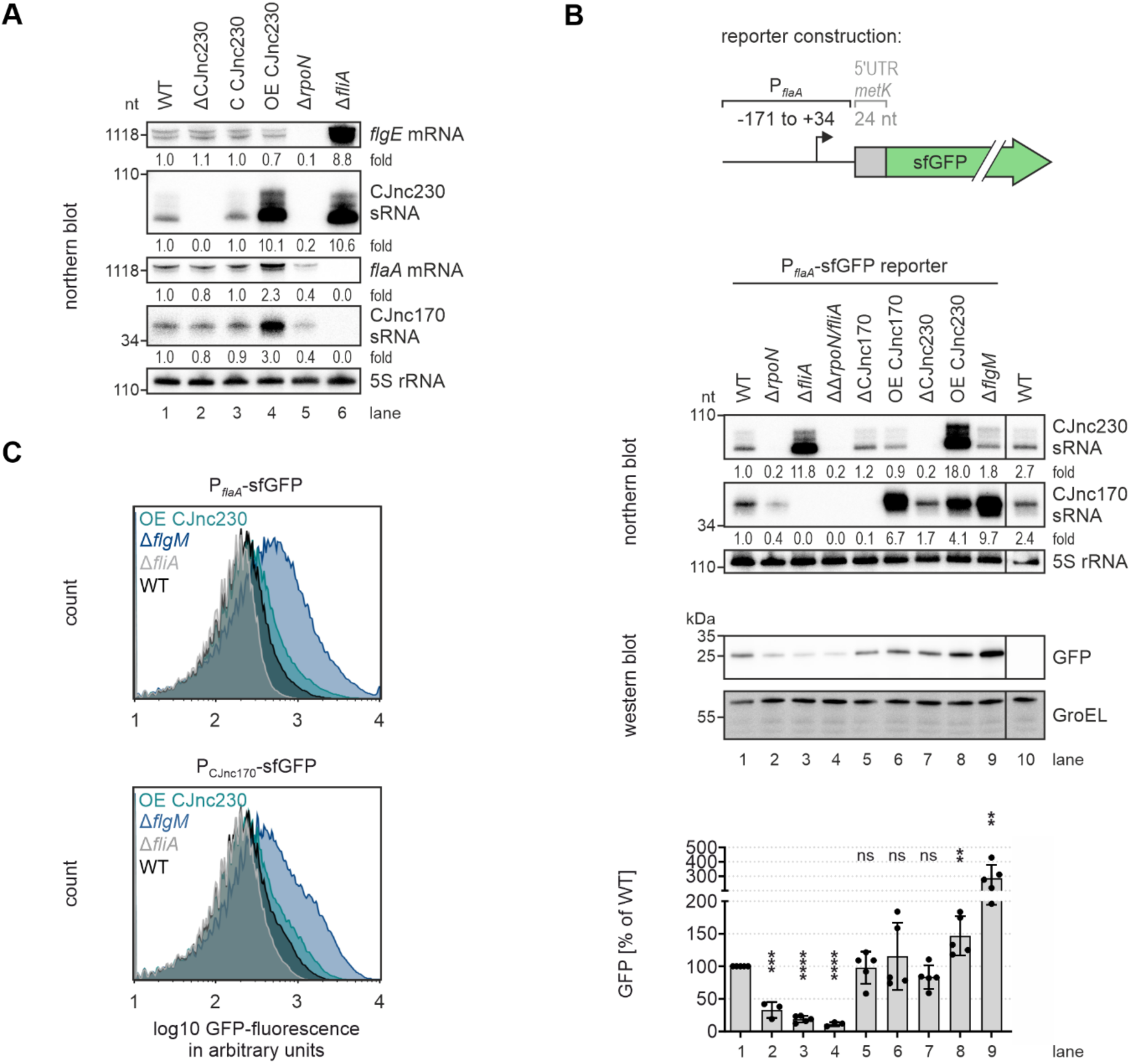
CJnc230-mediated post-transcriptional repression of *flgM* increases transcription of class III flagellar genes. **(A)** Northern blot analysis of total RNA from *C. jejuni* WT and CJnc230/sigma factor mutant strains grown to exponential phase. Transcripts were detected with the following radiolabeled oligonucleotides: *flgE* mRNA - CSO-5136, CJnc230 sRNA - CSO-0537, *flaA* mRNA - CSO-0486, CJnc170 sRNA - CSO-0182, 5S rRNA - CSO-0192 (loading control). Fold changes of sRNA/mRNA expression relative to WT and normalized to 5S rRNA are indicated. **(B)** Northern and western blots of *C. jejuni* WT, sigma factor, sRNA, or anti-sigma factor mutant strains with a transcriptional reporter of the *flaA* promoter region (nucleotide positions with respect to the *flaA* mRNA TSS (Dugar et al., 2013; Porcelli et al., 2013) are indicated in the scheme on top) fused to an unrelated ribosome binding site (RBS) (*metK*) and *sfgfp*, harvested at exponential growth phase. (*Middle panel*) CJnc230 and CJnc170 sRNA expression was probed on the northern blot with CSO-0537 and CSO-0182, respectively. 5S rRNA (CSO-0192): loading control. Fold changes of sRNA expression relative to the WT reporter and normalized to 5S rRNA are indicated. Representative images were cut between lanes 9 and 10. (*Lower panel*) Reporter expression measured by western blotting. GroEL was detected for normalization. One representative blot (cut between lanes 9 and 10) is shown with mean of independent replicates plotted below (n = 5 for WT, Δ*fliA*, ΔCJnc170, OE CJnc170, ΔCJnc230, OE CJnc230, and Δ*flgM*; n = 3 for Δ*rpoN* and Δ*rpoN*/Δ*fliA*). Error bars represent the standard deviation. ****: *p* < 0.0001, ***: *p* < 0.001, **: *p* < 0.01, ns: not significant, two-tailed Student’s *t*-test was used to compare the respective mutant to the WT reporter background (lane 1). **(C)** Flow cytometry analyses of P*flaA* (*upper*) and PCJnc170 (*lower*) transcriptional fusions grown to exponential phase. Representative histograms are shown. See also **Figure S9**.

To test whether increased *flaA* mRNA and CJnc170 sRNA levels observed upon CJnc230 overexpression arose from elevated transcription of class III promoters, we constructed transcriptional reporters including ∼200 nt upstream of their respective TSSs fused to an unrelated RBS (*metK*) and *sfgfp* (**Figs. 5B** **& S9A**). Deletion of *rpoN*, *fliA*, or *flgM* confirmed the functionality of the reporters, as expression measured on western blots and via flow cytometry was strongly reduced in Δ*rpoN* and Δ*fliA* backgrounds, and increased in Δ*flgM* (**Figs. 5B, 5C** & **S9**). Deletion or overexpression of CJnc230 slightly reduced or significantly upregulated P*flaA* reporter expression, respectively, by western blot analysis, compared to WT (**Fig. 5B**). In contrast, deletion or overexpression (∼7-fold, via the *porA* promoter) of FliA-dependent CJnc170 did not significantly affect *flaA* reporter expression. The significant upregulation of P*flaA*-sfGFP in OE CJnc230 was validated by flow cytometry, which further showed a significant reduction in P*flaA* reporter expression upon deletion of CJnc230 (**Figs. 5C** **& S9B**). Similar trends as for P*flaA*-sfGFP were observed for the PCJnc170 reporter. Overexpression of CJnc230 increased PCJnc170-sfGFP expression, but not quite to the levels of *flgM* deletion (**Figs. 5C** **& S9**).

Overall, the transcriptional reporter gene analyses indicate overexpression of CJnc230 increases transcript levels of class III genes of the flagellar regulatory cascade, *e.g.*, *flaA* and CJnc170. Moreover, our data show that this effect is due to increased FliA-dependent promoter activity, which is likely mediated by CJnc230 repression of the anti-σ^28^ factor FlgM.

### CJnc170 and CJnc230 balance flagellar biosynthesis and motility

So far, we have established a model for CJnc230 function in *C. jejuni* flagellar biogenesis in which repression of the anti-σ^28^ factor FlgM transcriptionally activates FliA-dependent class III flagellar genes such as the major flagellin *flaA* mRNA. This also upregulates the CJnc170 sRNA. CJnc170, together with its paralog CJnc10, was previously suggested to regulate *C. jejuni* class II flagellar genes, including *flgE* upstream of the CJnc230 sRNA (Le et al., 2015). Although no motility or flagellar morphology phenotype had been described so far for CJnc10 and CJnc170 deletion or overexpression mutants, the predicted targets of these two sRNAs point to possible feedback regulation on flagellar biosynthesis (Le et al., 2015). Since levels of CJnc10 seemed to be very low throughout growth in NCTC11168 (Dugar et al., 2013; Le et al., 2015) but CJnc170 was readily detectable on northern blot (**Figs. 1B** **& 5A**), we focused on CJnc170 for further experiments.

To explore a possible interplay between CJnc170 and CJnc230 in controlling *C. jejuni* motility, we constructed various combinations of sRNA single or double mutants. Above, we observed that overexpression, but not deletion, of CJnc230 in *C. jejuni* strain NCTC11168 significantly increased bacterial motility (**Figs. 4B** **& S10A**). In contrast, 21-fold overexpression of CJnc170 led to significantly reduced swimming motility compared to the WT (13.7 mm for WT vs. 8.2 mm for OE CJnc170), while ΔCJnc170 had no effect (**Figs. 6A** **& S10B**). Strikingly, deletion of CJnc230 in combination with CJnc170 overexpression completely abolished bacterial motility to levels observed for a Δ*flaA* mutant, revealing a phenotype not observed with single sRNA mutants before. In contrast, both deletion or overexpression of CJnc170 restored the increased motility phenotype of OE CJnc230 alone back to WT. Finally, the ΔCJnc170/ΔCJnc230 double deletion mutant showed similar, albeit significantly different, motility compared to WT (**Figs. 6A** **& S10B**).

**Figure 6.**
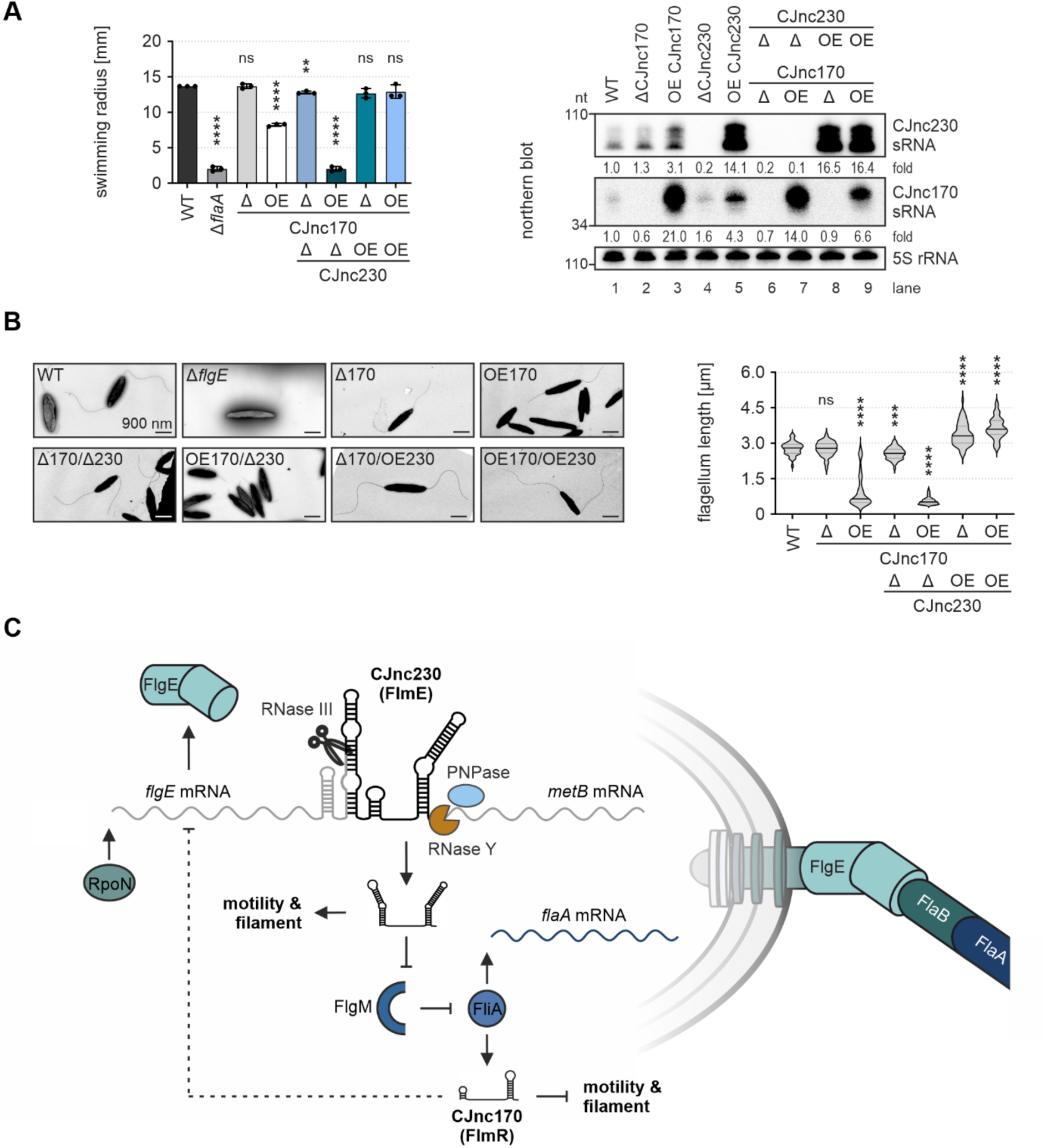
RpoN-dependent CJnc230 and FliA-dependent CJnc170 have opposing effects on motility and flagellar biogenesis. **(A)** (*Left*) Motility assays of *C. jejuni* WT and CJnc170 deletion (Δ) or overexpression (OE) mutants with or without deletion or overexpression of CJnc230 in 0.4% soft agar BB plates. Δ*flaA*: non-motile control. Error bars: standard deviation of the mean of independent replicates (n = 3). ****: *p* < 0.0001, **: *p* < 0.01, ns: not significant, two-tailed Student’s *t*-test vs. WT. A representative plate image is shown in **Figure S10B**. (*Right*) Northern blot analysis of total RNA from *C. jejuni* WT and CJnc170/CJnc230 deletion or OE mutants with or without deletion or overexpression of the respective other sRNA, harvested at exponential growth phase. CJnc230 and CJnc170 sRNA expression was probed with CSO-0537 and CSO-0182, respectively. 5S rRNA (CSO-0192): loading control. Fold changes of sRNA expression relative to WT and normalized to 5S rRNA are indicated. **(B)** Transmission electron microscopy (TEM) of *C. jejuni* WT, Δ*flgE*, and CJnc170/CJnc230 single or double deletion and/or OE mutants. (*Left*) Representative TEM images at 6,000 x magnification, scale bar = 900 nm. (*Right*) Violin plots depict the median (black) and quartiles (gray) of at least 44 flagellum length measurements per strain. Flagella are completely absent on Δ*flgE* bacteria and thus could not be quantified. ****: *p* < 0.0001, ***: *p* < 0.001, ns: not significant, two-tailed Student’s *t*-test was used to compare the respective mutant to the WT. **(C)** Model for CJnc230 (FlmE) and CJnc170 (FlmR) interplay in *C. jejuni* strain NCTC11168. CJnc230 is primarily transcribed together with the RpoN-dependent flagellar hook gene *flgE* and then processed downstream of the *flgE* CDS by RNase III. The sRNA 3’ end is further matured by RNase Y and PNPase. Overexpression of full-length CJnc230 (FlmE, 98 nt) increases filament length and motility, potentially via repression of the anti-σ^28^ factor FlgM, which de-represses class III flagellar genes such as *flaA* mRNA and the CJnc170 sRNA (dependent on FliA, σ^28^). Overexpression of CJnc170 (FlmR), in contrast, decreases motility and flagellar length, possibly via repression of class II flagellar genes such as *flgE* (Le et al., 2015) (dashed line). Overall, two hierarchically controlled sRNAs serve a regulatory circuit to control *C. jejuni* flagellar biogenesis.

Transmission electron microscopy analyses revealed that flagellar length strongly correlated with motility of the respective mutant (**Fig. 6B** **& Table S4**). We observed shorter filament length in the CJnc170 overexpression mutant, which might account for its reduced motility. However, significantly longer flagella were still found in CJnc230 overexpression mutants combined with CJnc170 deletion or overexpression (WT: 2,776 nm; ΔCJnc170: 2,765 nm; OE CJnc170: 972 nm; ΔCJnc170/ΔCJnc230: 2,566 nm; OE CJnc170/ΔCJnc230: 555 nm; ΔCJnc170/OE CJnc230: 3,373 nm; OE CJnc170/OE CJnc230: 3,640 nm) (**Fig. 6B** **& Table S4**). This suggests sRNA effects on motility which are independent of filament length and driven by yet unknown CJnc230 targets/functions. Based on their effects on filament length and bacterial motility, we propose renaming CJnc230/CJnc170 to FlmE/FlmR (flagellar length and <u>m</u>otility <u>e</u>nhancer/<u>r</u>epressor). In summary, CJnc230 and CJnc170 sRNAs have opposite effects on bacterial motility and filament assembly when heterologously overexpressed and uncoupled from native transcriptional regulation, suggesting a regulatory circuit in which the two sRNAs balance each other’s impact on flagellar biosynthesis (summarized in **Fig. 6C**).

## Discussion

In this study we have characterized a pair of *C. jejuni* sRNAs that are themselves hierarchically regulated by two flagellar sigma factors and have opposing effects on flagellar biosynthesis. CJnc230 (FlmE), processed from the RpoN-dependent *flgE* mRNA by RNase III, appears to upregulate class III flagellar gene expression via translational repression of the anti-σ^28^ factor FlgM. This increases FlaA levels, as well as flagellar length and motility. CJnc230 also influences expression of the class III sRNA CJnc170 (FlmR). In contrast to FlmE, FlmR reduces flagellar length and motility, possibly via feedback on class II genes, including *flgE* upstream of FlmE (Le et al., 2015). Together, our data point to an additional layer of post-transcriptional regulation in *C. jejuni* flagellar biosynthesis mediated by sRNAs, whose role might be to fine-tune the cascade. Together with the RpoN and FliA sigma factors, FlmE and FlmR comprise a mixed regulatory circuit, integrating transcriptional and post-transcriptional control (Brosse and Guillier, 2018), and possibly contribute to a faster switch from middle to late flagellar gene expression at reduced metabolic cost compared to protein factors (Nitzan et al., 2017)(**Fig. 6C**).

Reminiscent of FlmE and FlmR, several UTR-derived, σ^28^-dependent *E. coli* sRNAs have recently been described to have diverse effects on flagellin protein levels, flagellar number, and motility via regulation of middle and late flagellar genes, a transcriptional regulator, and a ribosomal protein mRNA (Melamed et al., 2021). Several other *E. coli* sRNAs post-transcriptionally regulate flagellar biosynthesis, but are controlled by environmental signals rather than flagellar sigma factors (*e.g*., EnvZ/OmpR and osmotic stress for OmrA/B) (Guillier and Gottesman, 2006; Prüß, 2017). Also, OmrA/B, ArcZ, OxyS, and McaS modulate the master regulator FlhDC (De Lay and Gottesman, 2012; Mika and Hengge, 2013; Thomason et al., 2012), which is absent in *C. jejuni*. Abundance of *flhD* mRNA also seems to be modulated by a *cis*-encoded RNA in conjunction with RNase III (Lejars et al., 2022). Only a handful of *E. coli* sRNAs target later points in the cascade. Like FlmE, *E. coli* OmrA/B sRNAs were recently shown to activate class III genes via repression of *flgM* mRNA (Romilly et al., 2020), while the Esr41 sRNA of enterohemorrhagic *E. coli* activates *fliA* transcription (Sudo et al., 2018, 2014; Waters et al., 2017).

FlmE, initially discovered during primary transcriptome analyses of *C. jejuni* (Dugar et al., 2013; Porcelli et al., 2013), caught our attention while looking for sRNAs that might be controlled by flagellar regulators. Our in-depth 5’-end mapping and co-transcriptional analyses indicate that a minor fraction is transcribed from its own RpoD-dependent promoter, while most of FlmE expression seems to be generated by processing from the same RpoN-dependent transcript as *flgE*, encoding the flagellar hook protein. FlmE adds to an expanding number of bacterial sRNAs that are processed by RNase III (Faubladier et al., 1990; Lalaouna et al., 2019; Melamed et al., 2020; Svensson and Sharma, 2021). So far, examples are mainly restricted to bacteria that lack RNase E, such as Gram-positive *Staphylococcus aureus* (RsaC, via cleavage of the *mntABC* mRNA 3’UTR (Lalaouna et al., 2019; Lioliou et al., 2012)) and *C. jejuni* (processing of the antisense sRNA pair CJnc180/190 (Svensson and Sharma, 2021)). While RNase III is required to generate the mature 5’ end of *flgE*-derived FlmE (**Fig. 2B**), the exact mechanism remains unclear. RNase III potentially cleaves a predicted intramolecular stem-loop downstream of the *flgE* CDS (**Fig. S3**), and maturation of the 5’ end might additionally involve trimming by exonucleases such as RNase J, which is essential in *C. jejuni* NCTC11168 (de Vries et al., 2017), or yet unidentified RNases as in the case of the RpoD-dependent FlmE precursor (**Fig. S5B, C**). *S. aureus* RNases J1/J2, for example, trim the 5’ end of *uhpT* mRNA to generate the 3’UTR-derived sRNA RsaG (Desgranges et al., 2022), and *Bacillus subtilis* RNase J1 trims the signal recognition particle (SRP) RNA component scRNA (small cytoplasmic RNA) after processing by RNase III (Yao et al., 2007). While 3’UTR-derived sRNAs control multiple physiological processes in diverse bacterial species (Miyakoshi et al., 2015; Ponath et al., 2022), also with implications for larger regulatory networks such as the bacterial envelope stress response (Chao and Vogel, 2016; Papenfort and Melamed, 2023), processing of flagellar mRNAs to produce sRNAs that function in the same pathway is a relatively new concept. To the best of our knowledge, only two sRNAs that potentially repress flagellar biogenesis and are also processed from flagellar mRNAs have been reported in *E. coli*: FlgO and FliX, which are processed from the 3’UTRs of FliA-dependent *flgL* and *fliC* mRNAs, respectively (Melamed et al., 2021; Thomason et al., 2015). While FlgO has a minor impact on flagellar number and motility, FliX seems to repress these two traits via regulation of several middle and late flagellar genes and an mRNA encoding ribosomal proteins (Melamed et al., 2021).

RNase Y and PNPase likely play a role in 3’ end maturation/degradation of FlmE (**Fig. 2A**). In contrast to the complete absence of mature FlmE sRNA in Δ*rnc*, these RNases only affected the abundance of the most stable 88-nt species (Δ*pnp*) or additional higher molecular weight fragments (Δ*rny* and Δ*rny*/Δ*pnp*). This hints at a combined action of endo- and exonucleases in *C. jejuni* RNA maturation or degradation as observed in *Streptococcus pyogenes* or *B. subtilis* (Broglia et al., 2020; Taggart et al., 2023). In Gram-positive bacteria such as *S. aureus* and *B. subtilis*, where RNase Y is highly conserved in place of RNase E (Bechhofer and Deutscher, 2019; Durand and Condon, 2018), it has global effects on the transcriptome, with 10-30% of genes differentially expressed upon its absence (Durand et al., 2012; Khemici et al., 2015; Laalami et al., 2013; Lehnik-Habrink et al., 2011). Our data suggest that RNase Y might have similar effects in Gram-negative *C. jejuni*, which likewise lacks RNase E, as approximately 5% of genes were affected in Δ*rny* (**Table S5**). This points to a possible broad regulatory role for *C. jejuni* RNase Y in transcriptome stability and sRNA processing.

Overexpression of FlmE increases motility, and we have identified two mRNA targets repressed by FlmE that might underlie this phenotype: Cj1387c and *flgM*. Cj1387c (CfiR, *Campylobacter* flagella interaction regulator) encodes a putative transcriptional regulator with a PAS domain that was reported to modulate flagella-flagella interactions via post-translational flagellar modification, leading to reduced autoagglutination of ΔCj1387c bacteria (Reuter et al., 2015). In *C. jejuni* strain 81-176, the Cj1387c ortholog (HeuR, heme uptake regulator) promotes chicken colonization and regulates iron metabolism and methionine biosynthesis, possibly via direct binding to promoter regions (Johnson et al., 2016; Kelley et al., 2021). This points to an additional role for FlmE not only in regulating flagellar biosynthesis, but also in important metabolic pathways via repression of Cj1387c, thus connecting these infection-relevant processes (Crofts et al., 2018; Liu et al., 2018; Ruddell et al., 2020).

The second validated target of FlmE repression is the anti-σ^28^ factor FlgM, a key hub in the checkpoint between class II and class III genes. In line with the role of FlgM in sequestering FliA activity (Wösten et al., 2010), we measured increased FlaA levels upon overexpression of FlmE (**Figs. 4A** **& 5A**). We also observed longer flagella in the FlmE overexpression strain, suggesting that an increase in FlaA levels might increase filament length (**Fig. 4C-E**). The *C. jejuni* filament length also impacts the secretion of virulence-associated factors (Barrero-Tobon and Hendrixson, 2014). Identification of conditions or additional regulators controlling FlmE and FlmR, or additional targets outside of flagella regulation like HeuR, might point to a role for these sRNAs beyond fine-tuning the biogenesis cascade to modulating flagellar morphology in response to different environments. This would expand the complexity of the system, whose temporal and spatial dynamics in individual bacteria might be disentangled by future studies with single-cell transcriptomics or imaging (Dar et al., 2021; Endesfelder, 2019; Homberger et al., 2022).

In *C. jejuni*, *flgM* is both co-transcribed with the upstream RpoN-dependent genes *flgI* and *flgJ*, as well as independently transcribed from its own FliA promoter (Dugar et al., 2013; Porcelli et al., 2013; Wösten et al., 2004). Temperature also affects *flgM* transcription and the interaction of FlgM protein with FliA (Stintzi, 2003; Wösten et al., 2010). Here, we further expand the regulatory complexity of the FlgM-FliA checkpoint inhibition strategy in *C. jejuni* with a layer of post-transcriptional control mediated by RpoN-dependent FlmE. The timing of regulation under native conditions, with *flgM* transcribed from two distinct promoters, is still unclear. This layer also likely interacts with the CsrA-FliW partner-switch mechanism that controls *flaA* mRNA translation and localization in *C. jejuni* (Dugar et al., 2016). This mechanism is distinct from Gram-negatives such as *E. coli*, where CsrA is titrated by sRNAs CsrB/C and activates the motility cascade through binding to *flhDC* mRNA (Wei et al., 2001), and more closely reflects Gram-positives. Together with the unique architecture of transcriptional control (*e.g.*, lack of the master regulator FlhDC) (Gilbreath et al., 2011; Lertsethtakarn et al., 2011), our results further set *C. jejuni* and the Epsilonproteobacteria apart as distinct models for flagella regulation.

Overall, our study establishes regulatory RNAs as key post-transcriptional modulators of *C. jejuni* flagellar filament assembly and motility. In the absence of a dedicated master regulator of flagellar gene expression, *C. jejuni* has evolved sophisticated transcriptional mechanisms to grant proper biogenesis of its flagellar machine, with alternative sigma factors RpoN and FliA at the forefront. Our presented work expands the regulatory complexity of the system by sRNA-mediated control of the hierarchical cascade, which fine-tunes *C. jejuni* motility, an important virulence trait of this human pathogen.

## Materials and methods

### Bacterial strains, plasmids, and oligonucleotides

Bacterial strains used in this study are listed in **Table S6**, while plasmids are in **Table S7**. DNA oligonucleotides used for cloning, T7 transcription template generation, and as northern blot probes are listed in **Table S8**.

### Bacterial standard growth

*E. coli* strains were grown aerobically at 37°C in Luria-Bertani (LB) broth or on LB agar, where necessary supplemented with 100 µg/ml ampicillin (Amp), 20 µg/ml chloramphenicol (Cm), 20 µg/ml gentamicin (Gm), or 20 µg/ml kanamycin (Kan) for marker selection. *E. coli* strains were stored at -80°C in LB media containing 10% DMSO.

*C. jejuni* was grown on Mueller-Hinton (MH) agar plates supplemented with 10 µg/ml vancomycin. For selection of transformants and growth of mutant strains, 50 µg/ml Kan, 20 µg/ml Cm, 20 µg/ml Gm, or 250 µg/ml hygromycin B (Hyg) were added to the plates. For liquid cultures, 10 ml or 15 ml Brucella broth (BB) medium supplemented with 10 µg/ml vancomycin was inoculated with *Campylobacter* cells from plate to a final OD600 nm of 0.002-0.005. Cultures were grown in 25 cm³ cell culture flasks shaking at 140 rpm until bacteria reached exponential/mid-log phase (OD600 nm of 0.4-0.5), or back-diluted to a final OD600 nm of 0.05 in 50 ml fresh BB media (75 cm³ flasks) for further growth as indicated in the text or figure legends. Plates and liquid cultures were incubated at 37°C in a HERAcell 150i incubator (Thermo Fisher Scientific) providing a microaerobic atmosphere (10% CO2, 5% O2, 85% N2). *C. jejuni* strains were stored at - 80°C in BB media containing 25% glycerol.

### Construction of *C. jejuni* deletion mutants

Deletion mutants of *C. jejuni* used in this study were constructed by double-crossover homologous recombination replacing the respective genomic region by an antibiotic resistance cassette. Cassettes used for cloning were either *aphA-3* (Kan^R^) (Skouloubris et al., 1998), *C. coli cat* (Cm^R^) (Wang and Taylor, 1990), *aac(3)-IV* (Gm^R^) (Bury-Moné et al., 2003), or *aph(7’’)* (Hyg^R^) (Cameron and Gaynor, 2014). By overlap PCR, these resistance cassettes were fused to ∼500 bp homologous sequences up- and downstream of the gene intended to be deleted.

As an example, deletion of the CJnc230 sRNA in *C. jejuni* strain NCTC11168 (CSS-5604) is described in detail. First, approximately 500 bp upstream of the CJnc230 5’ end and ∼500 bp downstream of the sRNA were amplified from NCTC11168 WT genomic DNA (gDNA) using CSO-3988 x 3989 and CSO-3991 x 3993, respectively. A polar Kan^R^ cassette (*aphA-3*) was amplified from pGG1 (Dugar et al., 2016) using JVO-5068 x HPK2-term. This cassette contained the promoter and terminator sequence from the *H. pylori* sRNA RepG (Pernitzsch et al., 2014). The antisense oligonucleotide (CSO-3989) for amplification of the CJnc230 upstream region and the sense oligonucleotide (CSO-3991) for amplification of the downstream region included overhangs (32 nt and 21 nt, respectively) to the resistance cassette in order to fuse all three fragments together by overlap PCR. Therefore, the purified PCR amplicons (Macherey-Nagel NucleoSpin Gel and PCR Clean-up Kit) of CJnc230 up- and downstream regions, as well as the polar Kan^R^ cassette were mixed in a 20:20:200 ng ratio and amplified using CSO-3988 x 3993 (final concentration 60 nM). After purification, the product was transformed into *C. jejuni* NCTC11168 WT strain (CSS-5295) via electroporation (described below). The resulting CJnc230::*aphA-3* transformants were checked by colony PCR using CSO-3995 x HPK2-term. A similar approach was chosen also for other deletion mutants used in this study, except for the use of non-polar resistance cassettes when replacing CDSs instead of sRNAs, and oligonucleotides are listed in **Table S8**.

### Construction of *C. jejuni* sRNA complementation and overexpression strains

For complementation or overexpression of CJnc170 and CJnc230, a copy of the respective sRNA was either introduced into the *rdxA* (Cj1066) or the Cj0046 pseudogene locus of *C. jejuni*, both of which were previously used for heterologous gene expression in this bacterium (Kim et al., 2008; Ribardo et al., 2010). For complementation of CJnc230, the sRNA was fused with its 5’ end to the unrelated *metK* (Cj1096c) promoter, as the majority of CJnc230 expression originated from cleavage of the *flgE* mRNA. Strong overexpression of either CJnc170 or CJnc230 was achieved by fusing it to the promoter of the major outer membrane protein (*porA*, Cj1259). Constructs were generated via sub-cloning of existing plasmids (pSE59.1 and pST1.1 (Dugar et al., 2018), or pSSv54.3 (Svensson and Sharma, 2021)) in *E. coli* TOP10 using homologous regions of the respective *C. jejuni* loci and *C. coli cat* (Cm^R^) (Wang and Taylor, 1990), *aphA-3* (Kan^R^) (Skouloubris et al., 1998), or *aac(3)-IV* (Gm^R^) (Bury-Moné et al., 2003) cassettes with promoter and terminator.

As an example, construction of the *C. jejuni* NCTC11168 CJnc230 complementation strain (CSS-6172) is described in detail. The CJnc230 sRNA starting from its 5’ end until 124 bp after its annotated 3’ end (Dugar et al., 2013) was amplified with CSO-4254 x 4257 from NCTC11168 WT (CSS-5295) gDNA and subsequently digested with *Cla*I. The plasmid backbone containing homologous parts to *rdxA*, a Cm^R^ (*cat*) cassette, and the *metK* promoter was amplified by inverse PCR with CSO-0347 x 1956 from pSE59.1 (Dugar et al., 2018) and also *Cla*I digested. Vector and insert were ligated overnight at 16°C using T4 DNA ligase (New England Biolabs) and transformed into *E. coli* TOP10. Positive clones were confirmed by colony PCR using CSO-0644 x 4257 and sequencing (Microsynth). The resulting plasmid was named pFK5.4. The purified PCR product, amplified from pFK5.4 with CSO-2276 x 2277, was then transformed into *C. jejuni* NCTC11168 ΔCJnc230 (CSS-5604) via electroporation. Clones were verified by colony PCR using CSO-0644 x 0349 and sequencing. A similar strategy was used to construct CJnc230 or CJnc170 overexpression mutants and respective oligonucleotides can be found in **Table S8**. In order to combine existing insertions in the *rdxA* locus (*e.g.* translational reporters; described below) with sRNA overexpression, plasmids based on pSSv54.3 (Svensson and Sharma, 2021) with regions homologous to the Cj0046 pseudogene locus were constructed. Site-directed mutagenesis of the CJnc230 interaction region was performed by inverse PCR on pFK16.7 and *Dpn*I digestion using primers CSO-5216 x 5217, resulting in pFK37.1.

### Construction of 3xFLAG epitope-tagged proteins in *C. jejuni*

In order to measure protein expression *in vivo*, the C-terminus of a protein of interest was chromosomally tagged with a 3xFLAG epitope. This was achieved by overlap PCR and is briefly explained for *C. jejuni* NCTC11168 FlgM-3xFLAG strain (CSS-5798).

For homologous recombination, approximately 500 bp upstream of the *flgM* stop codon and a region containing roughly the last 80 bp and 400 bp downstream of *flgM* CDS including parts of Cj1465 were amplified from NCTC11168 WT (CSS-5295) gDNA using oligonucleotides CSO-4431 x 4432 and CSO-4433 x 4434, respectively. The *aac(3)-IV* cassette (Gm^R^) fused downstream to the 3xFLAG sequence was amplified with primers CSO-0065 x HPK2 from gDNA of PtmG-3xFLAG (CSS-1252) (Svensson and Sharma, 2021). Oligonucleotides CSO-4432 and CSO-4433 generated overhangs to the 3xFLAG tag 5’ end and the Gm resistance cassette 3’ end, respectively. All three purified fragments were mixed in an equimolar ratio and used for overlap PCR (CSO-4431 x 4434, final concentration 60 nM). The purified PCR product was electroporated into NCTC11168 WT (CSS-5295) and correct insertion in resulting clones was validated by colony PCR using oligonucleotides CSO-4430 x HPK2 and sequencing. Analogous approaches were chosen for C-terminal epitope-tagging of Cj1387c, FlaA, FlaB (Dugar et al., 2016), RpoN, and FliA. The 3xFLAG fused to a kanamycin resistance cassette was amplified from pGG1 (Dugar et al., 2016). Respective oligonucleotides for cloning of epitope-tagged strains can be found in **Table S8**.

### Construction of *C. jejuni* sfGFP reporter fusions

For the construction of translational and transcriptional sfGFP reporters in NCTC11168, the 5’UTR or promoter region of interest was fused to an unrelated promoter or 5’UTR (*metK*), respectively. These constructs were linked to *sfgfp* (Pédelacq et al., 2006) and inserted into the *rdxA* (Cj1066) or Cj0046 pseudogene locus. Reporter fusions were generated by cloning in *E. coli* TOP10 and/or overlap PCR.

As an example for the overlap PCR approach, construction of the *flgM* (Cj1464) 5’UTR-sfGFP translational fusion in *C. jejuni* NCTC11168 (CSS-6300) is explained in more detail. First, the *rdxA* upstream region for homologous recombination including the *C. coli cat* cassette (Cm^R^) and the *metK* promoter (P*metK*), as well as *sfgfp* (starting from its 2nd codon) with the *rdxA* downstream region were amplified from pKF1.1 using oligonucleotides CSO-2276 x 1956 and CSO-3279 x 2277, respectively. This plasmid contained the *sfgfp* sequence amplified with CSO-3279 x 3717 on pXG10-SF (Corcoran et al., 2012), which was ligated into pSE59.1 (Dugar et al., 2018) after backbone amplification with CSO-4207 x 0347 and *Cla*I digestion. Positive *E. coli* TOP10 clones were confirmed after transformation by colony PCR using CSO-0644 x 3270 and sequencing. Second, the *flgM* 5’UTR including the predicted interaction region with CJnc230 and its first 10 codons were amplified from NCTC11168 WT (CSS-5295) gDNA using primers CSO-4721 x 4722. These oligonucleotides created overhangs to the P*metK* 3’ end (CSO-4721) or to the *sfgfp* 5’ end (CSO-4722). The resulting products were mixed in an equimolar ratio and used for overlap PCR (CSO-2276 x 2277, final primer concentration 60 nM). The purified PCR product was electroporated into NCTC11168 WT (CSS-5295) and correct integration was confirmed by colony PCR using oligonucleotides CSO-0644 x 0349 and sequencing. A similar approach was followed in order to generate the Cj1387c 5’UTR-sfGFP translational fusion for CJnc230 interaction validation.

Transcriptional reporter fusions were generated by a combination of cloning in *E. coli* TOP10 and subsequent overlap PCR. For better understanding, construction of the *C. jejuni* NCTC11168 P*flaA*-sfGFP strain (CSS-7086) is explained. First, the promoter region of *flaA* (position -171 to +34 relative to TSS) was amplified with CSO-5246 x 5247 from NCTC11168 WT (CSS-5295) gDNA and subsequently digested with *Xma*I. The vector backbone containing flanking regions of the *rdxA* locus for homologous recombination, the *C. coli cat* cassette (Cm^R^), the *metK* 5’UTR (24 nt, including RBS), as well as *sfgfp* was amplified from pKF1.1 using oligonucleotides CSO-2989 x 5096 and similarly digested. Vector and insert were ligated overnight at 16°C using T4 DNA ligase and transformed into *E. coli* TOP10. Positive clones were confirmed by colony PCR using CSO-0644 x 3270 and sequencing. The resulting plasmid was named pFK38.15. Next, the region containing the *flaA* promoter, *metK* 5’UTR, and *sfgfp* was amplified from pFK38.15 using oligonucleotides CSO-5248 x 5100, thereby generating overhangs to the 3’ end of the Gm^R^ cassette terminator and the Cj0046 flanking region, respectively. The Cj0046 up- and downstream regions for homologous recombination including the Gm^R^ cassette were amplified from pSSv54.3 (Svensson and Sharma, 2021) with CSO-1402 x 0484 and CSO-2751 x 1405, respectively. All three fragments were mixed in an equimolar ratio and used for overlap PCR (CSO-1402 x 1405, final primer concentration 60 nM). The purified PCR product was electroporated into NCTC11168 WT (CSS-5295) and correct integration was confirmed by colony PCR using oligonucleotides CSO-0833 x 3197 and sequencing. A similar approach was used for cloning of transcriptional reporter fusions with CJnc170, CJnc230, and *flgE* promoters. Respective oligonucleotides for cloning of translational and transcriptional sfGFP reporters are listed in **Table S8** and reporter sequences in **Table S9**.

### Construction of the FlaA (S395C) mutant for flagellar labeling

For imaging of flagellar filaments upon maleimide staining (see below), a *C. jejuni* NCTC11168 mutant bearing a second copy of *flaA* with an introduced cysteine residue (S395C) was generated (CSS-7487). The *flaA* locus including its FliA-dependent promoter (Dugar et al., 2013) was amplified from *C. jejuni* NCTC11168 WT (CSS-5295) gDNA using oligonucleotides CSO-4744 x 4745 and ligated into pGEM-T Easy (Promega) according to the manufacturer’s recommendations. Site-directed mutagenesis was performed on the resulting plasmid pFK43.1 by inverse PCR with primers CSO-5344 x 5345 followed by *Dpn*I digestion in order to mutate serine at position 395 to cysteine (S395C). A similar residue was previously used for mutation and subsequent flagellar staining in the related *C. jejuni* strain 81-176 (Cohen et al., 2020), as this is assumed to be surface-exposed according to its glycosylation status (Ewing et al., 2009). After colony PCR and sequencing with primers REV and UNI-61, this plasmid was named pFK44.3 and served as template for amplification of the mutated *flaA* (S395C) locus using oligonucleotides CSO-4744 x 4745 for subsequent overlap PCR. The 5’ end of this amplicon had complementarity to the upstream fragment for overlap PCR, containing ∼500 bp of the *rdxA* locus and the *C. coli cat* (Cm^R^) cassette, while the 3’ end of the amplicon overlapped with the downstream fragment of the *rdxA* locus for homologous recombination. The up- and downstream fragments for overlap PCR were amplified from gDNA of the NCTC11168 *flaA* complementation mutant (CSS-6208) (Alzheimer et al., 2020) with primers CSO-0345 x 0573 and CSO-0347 x 0348, respectively. All three products were mixed in an equimolar ratio and used for overlap PCR (CSO-2276 x 2277, final primer concentration 60 nM). The purified PCR product was electroporated into NCTC11168 WT (CSS-5295) and correct integration was confirmed by colony PCR using oligonucleotides CSO-0643 x 0349 and sequencing.

### Transformation of *C. jejuni* by electroporation

All mutant strains used in this study were generated by double-crossover homologous recombination in the chromosome after introducing DNA via electroporation. Therefore, *C. jejuni* was streaked out from cryostocks on MH agar with vancomycin, passaged once or maximum twice, and bacterial cells were resuspended in cold electroporation solution (272 mM sucrose, 15% (w/v) glycerol). Bacteria were pelleted by centrifugation at 4°C and 6,500 x *g* for 5 min, and cells were washed twice with the same solution. Depending on the pellet size, cells were taken up in 100-300 µl of cold electroporation solution and 50 µl of this suspension was mixed on ice with 200-400 ng of purified PCR product. Subsequently, the cells were electroporated in a 1 mm gap cuvette (Cell Projects) at 2.5 kV, 200 Ω, and 25 μF for 4-6 ms on a Biorad MicroPulser™ instrument. Afterwards, the electroporation reaction was mixed with 100 µl fresh BB medium at room temperature and transferred onto a non-selective MH agar plate with vancomycin only, to allow recovery overnight at 37°C in the microaerobic incubator. On the next day, cells were restreaked on a MH agar plate containing the appropriate antibiotics for marker selection and incubated microaerobically at 37°C until colonies were visible (2-5 days). Clones were verified by colony PCR after gene deletion and, in addition, by sequencing after gene insertions in *rdxA* or Cj0046 loci and C-terminal 3xFLAG-tagging.

### Motility assay

*C. jejuni* was grown to mid-log phase (OD600 nm 0.4-0.5) and 1 µl of each bacterial culture was stab-inoculated into a soft agar plate (BB + 0.4% Difco agar). Plates were incubated right-side-up at 37°C for 24 hrs under microaerobic conditions and, afterwards, the halo radius was measured at three different positions using ImageJ (NIH, USA; v 1.53j), thus averaging the mean swimming distance for each strain on one plate. Motility assays were performed in three biological replicates (three independent cultures stab-inoculated into three different plates) and the average halo radius for each mutant strain was used to compare motility to WT bacteria. Statistical tests as indicated were performed with GraphPad Prism (GraphPad Software, CA, USA; v 9.2.0).

### Transmission electron microscopy and flagellar length measurement

*C. jejuni* NCTC11168 were harvested from plate in passage one, gently soaked in PBS and centrifuged for 3 min at 2,500 x *g*. Pellets were carefully resuspended in 500 µl 2.5% glutaraldehyde including 0.1 M cacodylate and fixed overnight at 4°C. After staining with 2% uranylacetate, bacteria were imaged on 300-mesh grids using a JEOL-2100 transmission electron microscope. Flagellar length measurements were performed with ImageJ software (NIH, USA; v 1.53j) and the ridge detection plug-in applying the following settings: line width: 12; high contrast: 150; low contrast: 50; sigma: 3.96; lower threshold: 0.34; upper threshold: 0.85; minimum line length: 100; maximum line length: 6,000. For every strain, at least 30 seemingly intact flagella attached to the bacterial cell body were measured. Statistical tests were performed with GraphPad Prism (GraphPad Software, CA, USA; v 9.2.0).

### Maleimide-staining of flagella and confocal microscopy

In order to visualize flagellar filaments by confocal microscopy, *C. jejuni* NCTC11168 strains bearing a cysteine residue in a second copy of the major flagellin (FlaA S395C; described above) were labeled with the DyLight™ 488 maleimide stain (Thermo Scientific), adapted from a previously published protocol (Cohen et al., 2020). Therefore, bacteria grown on MH agar plates (passage one) corresponding to an OD600 nm of ∼1 were gently resuspended in 1 ml PBS and 1 µl of the maleimide-conjugated dye (stock concentration: 10 µg/µl in DMF) was added. After incubation at 37°C for 30 min, cells were centrifuged for 3 min at 2,500 x *g* and the pellet was carefully resuspended in 1 ml of PBS containing 1 µM of the eBioscience™ Cell Proliferation Dye eFluor™ 670 (Thermo Fisher Scientific) to counterstain the cell body. Following incubation at 37°C for 10 min and subsequent centrifugation, cells were washed once in PBS and then fixed for 2 hrs at room temperature in 4% paraformaldehyde/PBS. After two washes with PBS, samples were stored light-protected in the same buffer at 4°C until imaging with a Leica TCS SP5 II laser scanning confocal microscope (Leica Microsystems) using sequential scanning mode.

### SDS-PAGE and western blotting

*C. jejuni* cells grown to mid-log phase (OD600 nm 0.4-0.5) were collected by centrifugation at 7,500 x *g* and 4°C for 5 min. Pellets were dissolved in 1 x protein loading buffer (62.5 mM Tris-HCl, pH 6.8, 100 mM DTT, 10% (v/v) glycerol, 2% (w/v) SDS, 0.01% (w/v) bromophenol blue) and boiled for 8 min shaking at 1,000 rpm and 95°C.

For total protein analysis, samples corresponding to 0.1 OD600 nm were separated on vertical SDS-polyacrylamide gels (10 or 12% PAA) and stained by PageBlue protein staining solution (Thermo Scientific). For western blot analysis, protein samples corresponding to an OD600 nm of 0.05-0.2 were separated by SDS-PAGE (12 or 15% PAA) and transferred to a nitrocellulose membrane (GE) by semidry blotting. Membranes were blocked for 1 h with 10% (w/v) milk powder in TBS-T (Tris-buffered saline-Tween-20) and incubated overnight with primary antibody (monoclonal anti-FLAG, 1:1000; Sigma-Aldrich, #F1804-1MG; or anti-GFP, 1:1000, Roche #11814460001 in 3% bovine serum albumin (BSA)/TBS-T) at 4°C. After three washes for 20 min with TBS-T, membranes were incubated for 1 h at room temperature with secondary antibody (anti-mouse IgG, 1:10,000 in 3% BSA/TBS-T; GE Healthcare, #RPN4201) linked to horseradish peroxidase. After washing, chemiluminescence was detected using ECL-reagent (2 ml solution A (0.1 M Tris-HCl pH 8.6 and 0.25 mg/ml luminol sodium salt) and 200 µl solution B (1.1 mg/ml para-hydroxycomaric acid in DMSO)) with 0.6 µl 30% H2O2 and imaged on a ImageQuant LAS-4000 device (GE). A monoclonal antibody against GroEL (1:10,000; Sigma-Aldrich, #G6532-5ML) with an anti-rabbit IgG (1:10,000; GE Healthcare, #RPN4301) secondary antibody was used for normalization after FLAG/GFP detection. Bands were quantified using AIDA Image Analysis Software (Raytest, Germany). Statistical tests as indicated were performed with GraphPad Prism (GraphPad Software, CA, USA; v 9.2.0).

### Flow cytometry

For single cell analysis of sfGFP reporters in *C. jejuni*, bacteria grown to mid-log phase (OD600 nm 0.4-0.5) were collected by centrifugation at 7,500 x *g* and 4°C for 5 min. Pellets were resuspended in 500 µl 4% paraformaldehyde/PBS and fixed overnight at 4°C. After two washes with PBS, cells were resuspended in PBS and 100,000 events per sample were measured on a BD Accuri™ C6 instrument, applying a lower cutoff of 2,000 for the forward scatter (FSC-H). Analysis was done using FlowJo software (FlowJo, OR, USA; v10) and statistical tests were performed with GraphPad Prism (GraphPad Software, CA, USA; v 9.2.0).

### RNA preparation from bacterial cells

Unless stated otherwise, *C. jejuni* was grown to mid-log phase (OD600 nm 0.4-0.5) and cells corresponding to an OD600 nm of ∼4 were mixed with 0.2 volumes stop mix (95% EtOH and 5% phenol, v/v), and immediately snap-frozen in liquid nitrogen. Frozen cell pellets stored at -80°C were thawed on ice, spun down for 20 min at 4,500 x *g* and 4°C, and resuspended in 600 µl Tris-EDTA buffer (pH 8.0) containing 0.5 mg/ml lysozyme and 1% SDS. Cells were lysed upon incubation at 64°C for 2 min and, afterwards, total RNA was extracted using the hot phenol method as described previously for the related Epsilonproteobacterium *Helicobacter pylori* (Sharma et al., 2010). After RNA isolation, residual DNA was removed by treatment with DNase I (Thermo Scientific) according to the manufacturer’s instructions.

### Northern blot analysis

For RNA expression analyses, 5-10 μg of total RNA in Gel Loading Buffer II (GLII, Ambion; 95% (v/v) formamide, 18 mM EDTA, and 0.025% (w/v) SDS, xylene cyanol, and bromophenol blue) was separated on 6% PAA/7 M urea denaturing gels in 1 x TBE buffer. Afterwards, RNA was transferred to Hybond-XL membranes (GE) by electroblotting and cross-linked to the membrane with UV light. Then, γ^32^P-ATP end-labeled DNA oligonucleotides in Roti Hybri-quick (Roth) were hybridized overnight at 42°C. After washing at 42°C for 20 min each in 5 x, 1 x, and 0.5 x SSC (saline-sodium citrate) + 0.1% SDS, the membrane was dried and exposed to a PhosphorImager screen. Screens were scanned using a Typhoon FLA-7000 Series PhosphorImager (GE). Bands were quantified using AIDA Image Analysis Software (Raytest, Germany).

### Primer extension analysis

For identification of CJnc230 sRNA 5’ ends, 4 µg of DNase I-treated RNA from *C. jejuni* NCTC11168 WT and mutant strains was used for primer extension. RNA was diluted in H2O to a total volume of 5.5 μl, denatured, and snap-cooled on ice. The 5’-radiolabeled DNA oligonucleotide complementary to CJnc230 (CSO-0537) was added and annealed by heating to 80°C and slow cooling to 42°C. Afterwards, 3.5 µl of master mix containing reverse transcriptase (RT) buffer, dNTPs (1 mM final concentration) and 20 U Maxima RT (Thermo Scientific) was added and RNA was reverse transcribed for 1 h at 50°C. Reactions were stopped with 10 μl GLII. A sequencing ladder was generated using the DNA Cycle Sequencing kit (Jena Bioscience) according to the manufacturer’s recommendations with the CJnc230 sRNA region amplified from NCTC11168 WT (CSS-5295) gDNA using oligonucleotides CSO-3995 x 3993. The radioactively labeled CSO-0537 was also used for ladder preparation and reactions were stopped with GLII. Samples were separated on 6% PAA/7 M urea sequencing gels, which were dried afterwards and exposed to a PhosphorImager screen. Screens were scanned using a Typhoon FLA-7000 Series PhosphorImager (GE).

### Reverse transcription-polymerase chain reaction (RT-PCR)

For determination of *flgE*-CJnc230 co-transcription, DNase I-digested RNA samples of *C. jejuni* NCTC11168 WT and RNase deletion strains were reverse transcribed using the High Capacity cDNA Reverse Transcription kit (Thermo Fisher Scientific). One microgram of RNA was denatured, snap-cooled on ice, and reactions were started with or without (+/-) MultiScribe™ RT using the following cycling conditions: 25°C, 10 min; 37°C, 2 hrs; 85°C, 5 min. Afterwards, PCR was performed using gene-specific oligonucleotides for CJnc230 sRNA (CSO-4254 x 5138), as well as primer sets for amplification of CJnc230 together with the upstream gene *flgE* or with the downstream gene *metB* (CSO-5342 x 5138 or CSO-4254 x 5341, respectively). All three genes together were amplified using oligonucleotides CSO-5342 x 5341, and CJnc230 with 5’- or 3’-overhangs using primers CSO-4255 x 5138 or CSO-4254 x 4297, respectively. PCR on *C. jejuni* NCTC11168 WT (CSS-5295) gDNA served as positive control. Amplicons were visualized on 1-2% agarose gels, dependent on the expected fragment lengths.

### *In-vitro* transcription and 5’ end-labeling of RNAs

DNA templates containing the T7 promoter sequence were generated by PCR with S7 Fusion DNA polymerase (MOBIDING) using oligonucleotides listed in **Table S8**. *In-vitro* transcription of RNAs by T7 RNA polymerase was carried out using the MEGAscript™ T7 kit (Thermo Fisher Scientific) according to the manufacturer’s instructions and RNA quality was checked by electrophoresis on 6 or 10% PAA/7 M urea gels. Afterwards, transcripts were dephosphorylated with Antarctic Phosphatase (NEB), 5’ end-labeled (γ^32^P) with polynucleotide kinase (PNK; Thermo Scientific), and purified by gel extraction as described previously (Papenfort et al., 2006). Sequences of the resulting T7 transcripts and DNA templates are listed in **Table S10**.

### In-line probing

In order to determine RNA structure and binding interactions, in-line probing assays (Regulski and Breaker, 2008) were carried out as described previously (Pernitzsch et al., 2014). Five prime end-labeled CJnc230 (0.2 pmol, 20 nM final concentration) in the absence or presence of 2, 20, or 200 nM of *flgM* or Cj1387c mRNA leaders was incubated in 1 x in-line probing buffer (50 mM Tris-HCl, pH 8.3, 20 mM MgCl2, and 100 mM KCl) for 40 hrs at room temperature to allow spontaneous cleavage. RNA ladders were prepared using alkaline hydrolysis buffer (OH ladder) or sequencing buffer (RNase T1 ladder) according to the manufacturer’s instructions (Ambion). Reactions were stopped on ice with an equal volume of 2 x colorless loading buffer (10 M urea and 1.5 mM EDTA, pH 8.0), separated on 10% PAA/7 M Urea sequencing gels, and products were visualized after drying and exposure to PhosphorImager screens using a Typhoon FLA-7000 Series PhosphorImager (GE).

### *In-vitro* translation assay

Reporter fusions of sRNA targets were *in-vitro* translated in the absence and presence of sRNA using the PURExpress^Ⓡ^ *In Vitro* Protein Synthesis kit (NEB). Four picomoles of *in-vitro* transcribed 5’UTR reporters (including the RBS/sRNA interaction site and the first 10 codons of *flgM*, Cj1387c, or *flaA* fused to *sfgfp*; **Table S10**) were denatured in the absence or presence of 20, 80, or 200 pmol of CJnc230 sRNA for 1 min at 95°C and cooled on ice for 5 min. A *flaA* reporter was used as a negative control. The mRNA-sRNA mixture was pre-incubated for 10 min at 37°C before addition of the kit components, and translation was performed at 37°C for 30 min. Reactions were stopped with an equal volume of 2 x protein loading buffer. Fifteen microlitres were loaded on 12% PAA gels and protein expression was analyzed by western blotting with an antibody against GFP as described above. After blotting, residual protein in the gel was stained with PageBlue (Thermo Scientific) as a loading control.

### RNA-RNA interaction predictions

Predictions were computed genome-wide with IntaRNA (Mann et al., 2017) version 3.2.0 (linking Vienna RNA package 2.4.14) using default parameters, except for seed size of 6 nt. The 3’-truncated (88 nt) and full-length (98 nt) version of *C. jejuni* NCTC11168 CJnc230 (Dugar et al., 2013) were used as input and NC_002163.1 as genome accession number for strain NCTC11168.

### Total RNA sequencing (RNA-seq)

*C. jejuni* NCTC11168 WT and *rny* deletion strains were grown in biological triplicates in BB media to mid-log phase and cells were harvested for RNA isolation and subsequent DNase I digestion. Ribosomal RNA depletion and cDNA library preparation were performed at Vertis Biotechnologie AG, Germany. The adapter ligation protocol for construction of cDNA libraries is briefly summarized. After ultrasound fragmentation, an adapter was ligated to RNA 3’ ends and reverse transcription was performed, followed by ligation of 5’ Illumina TruSeq sequencing adapters to the 3’ ends of the antisense cDNAs. The resulting cDNA was PCR-amplified, purified with the Agencourt AMPure XP kit (Beckman Coulter Genomics), and analyzed by capillary electrophoresis. Samples were sequenced on an Illumina NextSeq 500 platform in single-end mode.

After sequencing, demultiplexing of reads was performed using bcl2fastq (Illumina; version: 2.20.0.422). Reads were quality and adapter trimmed with Cutadapt (versions: 1.17 and 2.5) (Martin, 2011) using a cutoff Phred score of 20 in NextSeq mode. READemption (version: 0.4.5) (Förstner et al., 2014) was used to align all reads longer than 11 nt to the reference genome of *C. jejuni* NCTC11168 (NCBI accession number: NC_002163.1; Refseq assembly accession GCF_000009085.1) using segemehl (version: 0.2.0) (Hoffmann et al., 2009) with an accuracy cutoff of 95%. Coverage plots representing the number of mapped reads per nucleotide were also generated by READemption and normalized for sequencing depth. NCBI gene annotations were complemented with annotations of previously determined 5’UTRs and sRNAs (Dugar et al., 2013) as described previously (Froschauer et al., 2022) and READemption was used to quantify aligned reads overlapping genomic features by at least 10 nt. Finally, differential gene expression analysis in replicates was performed by DESeq2 (version: 1.24.0) (Love et al., 2014) without fold-change shrinkage. Modeling of batch effects between experiments in the regression step was conducted by including a batch variable in the design formula. Genes with a log2FC ≥ |1| and an adjusted (Benjamini-Hochberg corrected) *p*-value ≤ 0.05 were considered as differentially expressed.

### Termination site sequencing (term-seq)

In order to determine transcript 3’ ends on a global scale in *C. jejuni* strain NCTC11168, we harvested WT and *rnc*, *rny*, *pnp*, and *rnr* mutant bacteria in biological duplicates for RNA isolation, followed by DNase I digestion. Ribosomal RNAs were depleted using the RiboCop™ rRNA Depletion Kit for Gram Negative Bacteria (Lexogen). Afterwards, cDNA libraries were constructed at Vertis Biotechnologie AG, Germany, using the 3’Term protocol. First, the 5’ Illumina sequencing adapter was ligated to the 3’-OH of the RNA molecules and cDNA synthesis was performed using M-MLV reverse transcriptase. Then, first-strand cDNA was fragmented and the 3’ Illumina sequencing adapter was ligated to the 3’ ends of the single-stranded cDNA fragments. Finally, the 3’ cDNA fragments were PCR-amplified and purified as described for total RNA-seq libraries above. Sequencing was performed using an Illumina NextSeq 500 platform in single-end mode.

Processing of sequencing reads, alignments, and coverage calculation were performed as described for total RNA-seq experiments. Then, the READemption pipeline (version: 0.4.5) (Förstner et al., 2014) was applied to generate two kinds of positional coverage files: default total coverage based on full-length alignments and last base coverage mapping of only the 3’-end base of each alignment. In both cases, sequencing depth-normalized plots were used for visualization in the Integrated Genome Browser (Freese et al., 2016).

### Conservation analyses and multiple sequence alignments

Putative CJnc230 homologs in different *Campylobacter* species were identified by blastn (Altschul et al., 1990), GLASSgo (Lott et al., 2018), and manual inspection of the intergenic region between Cj1729c (*flgE*) and Cj1727c (*metB*) homologs. Putative CJnc230 regions downstream of Cj1729c (*flgE*) homologs were retrieved from KEGG (Kyoto Encyclopedia of Genes and Genomes) and used for subsequent alignment with MultAlin (Corpet, 1988).

### Statistical analyses

All data for western blot, motility assay, flow cytometry, and growth behavior are presented as mean ± standard deviation. Flagellar length measurements are depicted as Violin plots including median and quartiles. Exact sample sizes of each experiment can be found in respective figure legends or in the main text. For statistical analysis, a two-tailed unpaired Student’s *t*-test was used. A value of *p* < 0.05 was considered significant and marked with an asterisk (*) as explained in the figure legends.

## Acknowledgements

We thank Gaurav Dugar and Sahil Sharma for help with cloning of ribonuclease deletion strains of *C. jejuni* NCTC11168, as well as the FlaB- and RpoN-3xFLAG strains. We are grateful to Gaurav Dugar for performing dRNA-seq experiments with NCTC11168 wild-type and Δ*rnc* strains and members of the Core Unit Systems Medicine (University of Würzburg and University Hospital Würzburg) for support with deep sequencing and bioinformatic analysis, as well as to Johannes Kullmann for mapping the dRNA-seq data. We thank the Imaging Core Facility at the Theodor-Boveri-Institute of Bioscience of the University of Würzburg for help with electron microscopy and Sharma lab members for critical comments and discussion on the manuscript.

## Author contributions

F.K. & C.M.S. designed the experiments; F.K. & S.L.S. performed the experiments; F.K., S.L.S. & C.M.S. analyzed data; F.K., S.L.S. & C.M.S. wrote the manuscript; C.M.S. provided resources.

## Main figures and legends

## References

1. Akahoshi DT, Bevins CL. 2022. Flagella at the Host-Microbe Interface: Key Functions Intersect With Redundant Responses. Front Immunol 13:828758. doi:10.3389/fimmu.2022.828758

2. Altschul SF, Gish W, Miller W, Myers EW, Lipman DJ. 1990. Basic local alignment search tool. J Mol Biol 215:403–410. doi:10.1016/S0022-2836(05)80360-2

3. Altuvia Y, Bar A, Reiss N, Karavani E, Argaman L, Margalit H. 2018. *In vivo* cleavage rules and target repertoire of RNase III in *Escherichia coli*. Nucleic Acids Res 46:10380–10394. doi:10.1093/nar/gky684

4. Alzheimer M, Svensson SL, König F, Schweinlin M, Metzger M, Walles H, Sharma CM. 2020. A three-dimensional intestinal tissue model reveals factors and small regulatory RNAs important for colonization with *Campylobacter jejuni*. PLoS Pathog 16:e1008304. doi:10.1371/journal.ppat.1008304

5. Balaban M, Joslin SN, Hendrixson DR. 2009. FlhF and its GTPase activity are required for distinct processes in flagellar gene regulation and biosynthesis in *Campylobacter jejuni*. J Bacteriol 191:6602–6611. doi:10.1128/JB.00884-09

6. Barrero-Tobon AM, Hendrixson DR. 2014. Flagellar biosynthesis exerts temporal regulation of secretion of specific *Campylobacter jejuni* colonization and virulence determinants. Mol Microbiol 93:957–974. doi:10.1111/mmi.12711

7. Barrero-Tobon AM, Hendrixson DR. 2012. Identification and analysis of flagellar coexpressed determinants (Feds) of *Campylobacter jejuni* involved in colonization. Mol Microbiol 84:352–369. doi:10.1111/j.1365-2958.2012.08027.x

8. Bechhofer DH, Deutscher MP. 2019. Bacterial ribonucleases and their roles in RNA metabolism. Crit Rev Biochem Mol Biol 54:242–300. doi:10.1080/10409238.2019.1651816

9. Broglia L, Lécrivain A-L, Renault TT, Hahnke K, Ahmed-Begrich R, Le Rhun A, Charpentier E. 2020. An RNA-seq based comparative approach reveals the transcriptome-wide interplay between 3’-to-5’ exoRNases and RNase Y. Nat Commun 11:1587. doi:10.1038/s41467-020-15387-6

10. Brosse A, Guillier M. 2018. Bacterial small RNAs in mixed regulatory networks. Microbiol Spectr 6. doi:10.1128/microbiolspec.RWR-0014-2017

11. Burnham PM, Hendrixson DR. 2018. *Campylobacter jejuni*: collective components promoting a successful enteric lifestyle. Nat Rev Microbiol 16:551–565. doi:10.1038/s41579-018-0037-9

12. Bury-Moné S, Skouloubris S, Dauga C, Thiberge J-M, Dailidiene D, Berg DE, Labigne A, De Reuse H. 2003. Presence of active aliphatic amidases in *Helicobacter* species able to colonize the stomach. Infect Immun 71:5613–5622. doi:10.1128/IAI.71.10.5613-5622.2003

13. Cameron A, Gaynor EC. 2014. Hygromycin B and apramycin antibiotic resistance cassettes for use in *Campylobacter jejuni*. PLoS ONE 9:e95084. doi:10.1371/journal.pone.0095084

14. Carrillo CD, Taboada E, Nash JHE, Lanthier P, Kelly J, Lau PC, Verhulp R, Mykytczuk O, Sy J, Findlay WA, Amoako K, Gomis S, Willson P, Austin JW, Potter A, Babiuk L, Allan B, Szymanski CM. 2004. Genome-wide expression analyses of *Campylobacter jejuni* NCTC11168 reveals coordinate regulation of motility and virulence by *flhA*. J Biol Chem 279:20327–20338. doi:10.1074/jbc.M401134200

15. Chao Y, Vogel J. 2016. A 3’ UTR-Derived Small RNA Provides the Regulatory Noncoding Arm of the Inner Membrane Stress Response. Mol Cell 61:352–363. doi:10.1016/j.molcel.2015.12.023

16. Chaudhuri RR, Yu L, Kanji A, Perkins TT, Gardner PP, Choudhary J, Maskell DJ, Grant AJ. 2011. Quantitative RNA-seq analysis of the *Campylobacter jejuni* transcriptome. *Microbiology (Reading*, Engl) 157:2922–2932. doi:10.1099/mic.0.050278-0

17. Chevance FFV, Hughes KT. 2008. Coordinating assembly of a bacterial macromolecular machine. Nat Rev Microbiol 6:455–465. doi:10.1038/nrmicro1887

18. Cohen EJ, Nakane D, Kabata Y, Hendrixson DR, Nishizaka T, Beeby M. 2020. *Campylobacter jejuni* motility integrates specialized cell shape, flagellar filament, and motor, to coordinate action of its opposed flagella. PLoS Pathog 16:e1008620. doi:10.1371/journal.ppat.1008620

19. Colin R, Ni B, Laganenka L, Sourjik V. 2021. Multiple functions of flagellar motility and chemotaxis in bacterial physiology. FEMS Microbiol Rev 45. doi:10.1093/femsre/fuab038

20. Corcoran CP, Podkaminski D, Papenfort K, Urban JH, Hinton JCD, Vogel J. 2012. Superfolder GFP reporters validate diverse new mRNA targets of the classic porin regulator, MicF RNA. Mol Microbiol 84:428–445. doi:10.1111/j.1365-2958.2012.08031.x

21. Corpet F. 1988. Multiple sequence alignment with hierarchical clustering. Nucleic Acids Res 16:10881–10890. doi:10.1093/nar/16.22.10881

22. Correa NE, Barker JR, Klose KE. 2004. The *Vibrio cholerae* FlgM homologue is an anti-sigma28 factor that is secreted through the sheathed polar flagellum. J Bacteriol 186:4613–4619. doi:10.1128/JB.186.14.4613-4619.2004

23. Crofts AA, Poly FM, Ewing CP, Kuroiwa JM, Rimmer JE, Harro C, Sack D, Talaat KR, Porter CK, Gutierrez RL, DeNearing B, Brubaker J, Laird RM, Maue AC, Jaep K, Alcala A, Tribble DR, Riddle MS, Ramakrishnan A, McCoy AJ, Trent MS. 2018. *Campylobacter jejuni* transcriptional and genetic adaptation during human infection. Nat Microbiol 3:494–502. doi:10.1038/s41564-018-0133-7

24. Dar D, Dar N, Cai L, Newman DK. 2021. Spatial transcriptomics of planktonic and sessile bacterial populations at single-cell resolution. Science 373:abi4882. doi:10.1126/science.abi4882

25. Dar D, Shamir M, Mellin JR, Koutero M, Stern-Ginossar N, Cossart P, Sorek R. 2016. Term-seq reveals abundant ribo-regulation of antibiotics resistance in bacteria. Science 352:aad9822. doi:10.1126/science.aad9822

26. Desgranges E, Barrientos L, Herrgott L, Marzi S, Toledo-Arana A, Moreau K, Vandenesch F, Romby P, Caldelari I. 2022. The 3’UTR-derived sRNA RsaG coordinates redox homeostasis and metabolism adaptation in response to glucose-6-phosphate uptake in *Staphylococcus aureus*. Mol Microbiol 117:193–214. doi:10.1111/mmi.14845

27. De Lay N, Gottesman S. 2012. A complex network of small non-coding RNAs regulate motility in *Escherichia coli*. Mol Microbiol 86:524–538. doi:10.1111/j.1365-2958.2012.08209.x

28. de Vries SP, Gupta S, Baig A, Wright E, Wedley A, Jensen AN, Lora LL, Humphrey S, Skovgård H, Macleod K, Pont E, Wolanska DP, L’Heureux J, Mobegi FM, Smith DGE, Everest P, Zomer A, Williams N, Wigley P, Humphrey T, Grant AJ. 2017. Genome-wide fitness analyses of the foodborne pathogen *Campylobacter jejuni* in *in vitro* and *in vivo* models. Sci Rep 7:1251. doi:10.1038/s41598-017-01133-4

29. Dugar G, Herbig A, Förstner KU, Heidrich N, Reinhardt R, Nieselt K, Sharma CM. 2013. High-resolution transcriptome maps reveal strain-specific regulatory features of multiple *Campylobacter jejuni* isolates. PLoS Genet 9:e1003495. doi:10.1371/journal.pgen.1003495

30. Dugar G, Leenay RT, Eisenbart SK, Bischler T, Aul BU, Beisel CL, Sharma CM. 2018. CRISPR RNA-Dependent Binding and Cleavage of Endogenous RNAs by the *Campylobacter jejuni* Cas9. Mol Cell 69:893–905.e7. doi:10.1016/j.molcel.2018.01.032

31. Dugar G, Svensson SL, Bischler T, Wäldchen S, Reinhardt R, Sauer M, Sharma CM. 2016. The CsrA-FliW network controls polar localization of the dual-function flagellin mRNA in *Campylobacter jejuni*. Nat Commun 7:11667. doi:10.1038/ncomms11667

32. Durand S, Condon C. 2018. RNases and Helicases in Gram-Positive Bacteria. Microbiol Spectr 6. doi:10.1128/microbiolspec.RWR-0003-2017

33. Durand S, Gilet L, Bessières P, Nicolas P, Condon C. 2012. Three essential ribonucleases-RNase Y, J1, and III-control the abundance of a majority of *Bacillus subtilis* mRNAs. PLoS Genet 8:e1002520. doi:10.1371/journal.pgen.1002520

34. Endesfelder U. 2019. From single bacterial cell imaging towards *in vivo* single-molecule biochemistry studies. Essays Biochem 63:187–196. doi:10.1042/EBC20190002

35. Ewing CP, Andreishcheva E, Guerry P. 2009. Functional characterization of flagellin glycosylation in *Campylobacter jejuni* 81-176. J Bacteriol 191:7086–7093. doi:10.1128/JB.00378-09

36. Faubladier M, Cam K, Bouché JP. 1990. *Escherichia coli* cell division inhibitor DicF-RNA of the *dicB* operon. Evidence for its generation *in vivo* by transcription termination and by RNase III and RNase E-dependent processing. J Mol Biol 212:461–471. doi:10.1016/0022-2836(90)90325-G

37. Förstner KU, Vogel J, Sharma CM. 2014. READemption-a tool for the computational analysis of deep-sequencing-based transcriptome data. Bioinformatics 30:3421–3423. doi:10.1093/bioinformatics/btu533

38. Freese NH, Norris DC, Loraine AE. 2016. Integrated genome browser: visual analytics platform for genomics. Bioinformatics 32:2089–2095. doi:10.1093/bioinformatics/btw069

39. Froschauer K, Svensson SL, Gelhausen R, Fiore E, Kible P, Klaude A, Kucklick M, Fuchs S, Eggenhofer F, Engelmann S, Backofen R, Sharma CM. 2022. Complementary Ribo-seq approaches map the translatome and provide a small protein census in the foodborne pathogen *Campylobacter jejuni*. BioRxiv. doi:10.1101/2022.11.09.515450

40. Gao B, Vorwerk H, Huber C, Lara-Tejero M, Mohr J, Goodman AL, Eisenreich W, Galán JE, Hofreuter D. 2017. Metabolic and fitness determinants for *in vitro* growth and intestinal colonization of the bacterial pathogen *Campylobacter jejuni*. PLoS Biol 15:e2001390. doi:10.1371/journal.pbio.2001390

41. Gilbreath JJ, Cody WL, Merrell DS, Hendrixson DR. 2011. Change is good: variations in common biological mechanisms in the epsilonproteobacterial genera *Campylobacte*r and *Helicobacter*. Microbiol Mol Biol Rev 75:84–132. doi:10.1128/MMBR.00035-10

42. Gillen KL, Hughes KT. 1991. Negative regulatory loci coupling flagellin synthesis to flagellar assembly in *Salmonella* typhimurium. J Bacteriol 173:2301–2310. doi:10.1128/jb.173.7.2301-2310.1991

43. Gruber AR, Lorenz R, Bernhart SH, Neuböck R, Hofacker IL. 2008. The Vienna RNA websuite. Nucleic Acids Res 36:W70–4. doi:10.1093/nar/gkn188

44. Guerry P. 2007. *Campylobacter* flagella: not just for motility. Trends Microbiol 15:456–461. doi:10.1016/j.tim.2007.09.006

45. Guillier M, Gottesman S. 2006. Remodelling of the *Escherichia coli* outer membrane by two small regulatory RNAs. Mol Microbiol 59:231–247. doi:10.1111/j.1365-2958.2005.04929.x

46. Haddad N, Matos RG, Pinto T, Rannou P, Cappelier J-M, Prévost H, Arraiano CM. 2014. The RNase R from *Campylobacter jejuni* has unique features and is involved in the first steps of infection. J Biol Chem 289:27814–27824. doi:10.1074/jbc.M114.561795

47. Haddad N, Saramago M, Matos RG, Prévost H, Arraiano CM. 2013. Characterization of the biochemical properties of *Campylobacter jejuni* RNase III. Biosci Rep 33. doi:10.1042/BSR20130090

48. Haiko J, Westerlund-Wikström B. 2013. The role of the bacterial flagellum in adhesion and virulence. Biology (Basel) 2:1242–1267. doi:10.3390/biology2041242

49. Havelaar AH, Kirk MD, Torgerson PR, Gibb HJ, Hald T, Lake RJ, Praet N, Bellinger DC, de Silva NR, Gargouri N, Speybroeck N, Cawthorne A, Mathers C, Stein C, Angulo FJ, Devleesschauwer B, World Health Organization Foodborne Disease Burden Epidemiology Reference Group. 2015. World health organization global estimates and regional comparisons of the burden of foodborne disease in 2010. PLoS Med 12:e1001923. doi:10.1371/journal.pmed.1001923

50. Hoffmann S, Otto C, Kurtz S, Sharma CM, Khaitovich P, Vogel J, Stadler PF, Hackermüller J. 2009. Fast mapping of short sequences with mismatches, insertions and deletions using index structures. PLoS Comput Biol 5:e1000502. doi:10.1371/journal.pcbi.1000502

51. Homberger C, Barquist L, Vogel J. 2022. Ushering in a new era of single-cell transcriptomics in bacteria. microLife 3:uqac020. doi:10.1093/femsml/uqac020

52. Hör J, Matera G, Vogel J, Gottesman S, Storz G. 2020. *Trans*-acting small RNAs and their effects on gene expression in *Escherichia coli* and *Salmonella enterica*. Ecosal Plus 9:ESP-0030-2019. doi:10.1128/ecosalplus.ESP-0030-2019

53. Janssen R, Krogfelt KA, Cawthraw SA, van Pelt W, Wagenaar JA, Owen RJ. 2008. Host-pathogen interactions in *Campylobacter* infections: the host perspective. Clin Microbiol Rev 21:505–518. doi:10.1128/CMR.00055-07

54. Johnson JG, Gaddy JA, DiRita VJ. 2016. The PAS Domain-Containing Protein HeuR Regulates Heme Uptake in *Campylobacter jejuni*. MBio 7. doi:10.1128/mBio.01691-16

55. Josenhans C, Niehus E, Amersbach S, Hörster A, Betz C, Drescher B, Hughes KT, Suerbaum S. 2002. Functional characterization of the antagonistic flagellar late regulators FliA and FlgM of *Helicobacter pylori* and their effects on the *H. pylori* transcriptome. Mol Microbiol 43:307–322. doi:10.1046/j.1365-2958.2002.02765.x

56. Joslin SN, Hendrixson DR. 2009. Activation of the *Campylobacter jejuni* FlgSR two-component system is linked to the flagellar export apparatus. J Bacteriol 191:2656–2667. doi:10.1128/JB.01689-08

57. Joslin SN, Hendrixson DR. 2008. Analysis of the *Campylobacter jejuni* FlgR response regulator suggests integration of diverse mechanisms to activate an NtrC-like protein. J Bacteriol 190:2422–2433. doi:10.1128/JB.01827-07

58. Kamal N, Dorrell N, Jagannathan A, Turner SM, Constantinidou C, Studholme DJ, Marsden G, Hinds J, Laing KG, Wren BW, Penn CW. 2007. Deletion of a previously uncharacterized flagellar-hook-length control gene *fliK* modulates the sigma54-dependent regulon in *Campylobacter jejuni*. Microbiology (Reading, Engl) 153:3099–3111. doi:10.1099/mic.0.2007/007401-0

59. Kelley BR, Callahan SM, Johnson JG. 2021. Transcription of Cystathionine β-Lyase (MetC) Is Repressed by HeuR in *Campylobacter jejuni*, and Methionine Biosynthesis Facilitates Colonocyte Invasion. J Bacteriol 203:e0016421. doi:10.1128/JB.00164-21

60. Khemici V, Prados J, Linder P, Redder P. 2015. Decay-Initiating Endoribonucleolytic Cleavage by RNase Y Is Kept under Tight Control via Sequence Preference and Sub-cellular Localisation. PLoS Genet 11:e1005577. doi:10.1371/journal.pgen.1005577

61. Kim J-S, Li J, Barnes IHA, Baltzegar DA, Pajaniappan M, Cullen TW, Trent MS, Burns CM, Thompson SA. 2008. Role of the *Campylobacter jejuni* Cj1461 DNA methyltransferase in regulating virulence characteristics. J Bacteriol 190:6524–6529. doi:10.1128/JB.00765-08

62. Kutsukake K, Iino T. 1994. Role of the FliA-FlgM regulatory system on the transcriptional control of the flagellar regulon and flagellar formation in *Salmonella* typhimurium. J Bacteriol 176:3598–3605. doi:10.1128/jb.176.12.3598-3605.1994

63. Laalami S, Bessières P, Rocca A, Zig L, Nicolas P, Putzer H. 2013. *Bacillus subtilis* RNase Y activity in vivo analysed by tiling microarrays. PLoS ONE 8:e54062. doi:10.1371/journal.pone.0054062

64. Lalaouna D, Baude J, Wu Z, Tomasini A, Chicher J, Marzi S, Vandenesch F, Romby P, Caldelari I, Moreau K. 2019. RsaC sRNA modulates the oxidative stress response of *Staphylococcus aureus* during manganese starvation. Nucleic Acids Res 47:9871–9887. doi:10.1093/nar/gkz728

65. Lejars M, Caillet J, Solchaga-Flores E, Guillier M, Plumbridge J, Hajnsdorf E. 2022. Regulatory Interplay between RNase III and Antisense RNAs in *E. coli*: the Case of AsflhD and FlhD, Component of the Master Regulator of Motility. MBio 13:e0098122. doi:10.1128/mbio.00981-22

66. Lertsethtakarn P, Ottemann KM, Hendrixson DR. 2011. Motility and chemotaxis in *Campylobacter* and *Helicobacter* . Annu Rev Microbiol 65:389–410. doi:10.1146/annurev-micro-090110-102908

67. Le MT, van Veldhuizen M, Porcelli I, Bongaerts RJ, Gaskin DJH, Pearson BM, van Vliet AHM. 2015. Conservation of σ28-Dependent Non-Coding RNA Paralogs and Predicted σ54-Dependent Targets in Thermophilic *Campylobacter* Species. PLoS ONE 10:e0141627. doi:10.1371/journal.pone.0141627

68. Lehnik-Habrink M, Schaffer M, Mäder U, Diethmaier C, Herzberg C, Stülke J. 2011. RNA processing in *Bacillus subtilis*: identification of targets of the essential RNase Y. Mol Microbiol 81:1459–1473. doi:10.1111/j.1365-2958.2011.07777.x

69. Lioliou E, Sharma CM, Caldelari I, Helfer A-C, Fechter P, Vandenesch F, Vogel J, Romby P. 2012. Global regulatory functions of the *Staphylococcus aureus* endoribonuclease III in gene expression. PLoS Genet 8:e1002782. doi:10.1371/journal.pgen.1002782

70. Liu MM, Boinett CJ, Chan ACK, Parkhill J, Murphy MEP, Gaynor EC. 2018. Investigating the *Campylobacter jejuni* Transcriptional Response to Host Intestinal Extracts Reveals the Involvement of a Widely Conserved Iron Uptake System. MBio 9. doi:10.1128/mBio.01347-18

71. Liu X, Gao B, Novik V, Galán JE. 2012. Quantitative Proteomics of Intracellular *Campylobacter jejuni* Reveals Metabolic Reprogramming. PLoS Pathog 8:e1002562. doi:10.1371/journal.ppat.1002562

72. Lott SC, Schäfer RA, Mann M, Backofen R, Hess WR, Voß B, Georg J. 2018. GLASSgo - Automated and Reliable Detection of sRNA Homologs From a Single Input Sequence. Front Genet 9:124. doi:10.3389/fgene.2018.00124

73. Love MI, Huber W, Anders S. 2014. Moderated estimation of fold change and dispersion for RNA-seq data with DESeq2. Genome Biol 15:550. doi:10.1186/s13059-014-0550-8

74. Mann M, Wright PR, Backofen R. 2017. IntaRNA 2.0: enhanced and customizable prediction of RNA-RNA interactions. Nucleic Acids Res 45:W435–W439. doi:10.1093/nar/gkx279

75. Martin M. 2011. Cutadapt removes adapter sequences from high-throughput sequencing reads. EMBnet j 17:10. doi:10.14806/ej.17.1.200

76. McSweegan E, Walker RI. 1986. Identification and characterization of two *Campylobacter jejuni* adhesins for cellular and mucous substrates. Infect Immun 53:141–148. doi:10.1128/iai.53.1.141-148.1986

77. Mediati DG, Wong JL, Gao W, McKellar S, Pang CNI, Wu S, Wu W, Sy B, Monk IR, Biazik JM, Wilkins MR, Howden BP, Stinear TP, Granneman S, Tree JJ. 2022. RNase III-CLASH of multi-drug resistant *Staphylococcus aureus* reveals a regulatory mRNA 3’UTR required for intermediate vancomycin resistance. Nat Commun 13:3558. doi:10.1038/s41467-022-31177-8

78. Melamed S, Adams PP, Zhang A, Zhang H, Storz G. 2020. RNA-RNA Interactomes of ProQ and Hfq Reveal Overlapping and Competing Roles. Mol Cell 77:411–425.e7. doi:10.1016/j.molcel.2019.10.022

79. Melamed S, Zhang A, Jarnik M, Mills J, Zhang H, Storz G. 2021. σ28-dependent small RNAs regulate timing of flagella biosynthesis. BioRxiv. doi:10.1101/2021.08.05.455139

80. Mika F, Hengge R. 2013. Small Regulatory RNAs in the Control of Motility and Biofilm Formation in *E. coli* and *Salmonella*. Int J Mol Sci 14:4560–4579. doi:10.3390/ijms14034560

81. Miyakoshi M, Chao Y, Vogel J. 2015. Regulatory small RNAs from the 3’ regions of bacterial mRNAs. Curr Opin Microbiol 24:132–139. doi:10.1016/j.mib.2015.01.013

82. Nitzan M, Rehani R, Margalit H. 2017. Integration of bacterial small RNAs in regulatory networks. Annu Rev Biophys 46:131–148. doi:10.1146/annurev-biophys-070816-034058

83. Papenfort K, Melamed S. 2023. Small RNAs, Large Networks: Posttranscriptional Regulons in Gram-Negative Bacteria. Annu Rev Microbiol. doi:10.1146/annurev-micro-041320-025836

84. Papenfort K, Pfeiffer V, Mika F, Lucchini S, Hinton JCD, Vogel J. 2006. SigmaE-dependent small RNAs of *Salmonella* respond to membrane stress by accelerating global *omp* mRNA decay. Mol Microbiol 62:1674–1688. doi:10.1111/j.1365-2958.2006.05524.x

85. Parkhill J, Wren BW, Mungall K, Ketley JM, Churcher C, Basham D, Chillingworth T, Davies RM, Feltwell T, Holroyd S, Jagels K, Karlyshev AV, Moule S, Pallen MJ, Penn CW, Quail MA, Rajandream MA, Rutherford KM, van Vliet AH, Whitehead S, Barrell BG. 2000. The genome sequence of the food-borne pathogen *Campylobacter jejuni* reveals hypervariable sequences. Nature 403:665–668. doi:10.1038/35001088

86. Pédelacq J-D, Cabantous S, Tran T, Terwilliger TC, Waldo GS. 2006. Engineering and characterization of a superfolder green fluorescent protein. Nat Biotechnol 24:79–88. doi:10.1038/nbt1172

87. Pernitzsch SR, Tirier SM, Beier D, Sharma CM. 2014. A variable homopolymeric G-repeat defines small RNA-mediated posttranscriptional regulation of a chemotaxis receptor in *Helicobacter pylori*. Proc Natl Acad Sci USA 111:E501–10. doi:10.1073/pnas.1315152111

88. Ponath F, Hör J, Vogel J. 2022. An overview of gene regulation in bacteria by small RNAs derived from mRNA 3’ ends. FEMS Microbiol Rev 46. doi:10.1093/femsre/fuac017

89. Porcelli I, Reuter M, Pearson BM, Wilhelm T, van Vliet AHM. 2013. Parallel evolution of genome structure and transcriptional landscape in the Epsilonproteobacteria. BMC Genomics 14:616. doi:10.1186/1471-2164-14-616

90. Prüß BM. 2017. Involvement of Two-Component Signaling on Bacterial Motility and Biofilm Development. J Bacteriol 199. doi:10.1128/JB.00259-17

91. Radomska KA, Wösten MMSM, Ordoñez SR, Wagenaar JA, van Putten JPM. 2017. Importance of *Campylobacter jejuni* FliS and FliW in Flagella Biogenesis and Flagellin Secretion. Front Microbiol 8:1060. doi:10.3389/fmicb.2017.01060

92. Regulski EE, Breaker RR. 2008. In-line probing analysis of riboswitches. Methods Mol Biol 419:53–67. doi:10.1007/978-1-59745-033-1_4

93. Reuter M, Periago PM, Mulholland F, Brown HL, van Vliet AHM. 2015. A PAS domain-containing regulator controls flagella-flagella interactions in *Campylobacter jejuni*. Front Microbiol 6:770. doi:10.3389/fmicb.2015.00770

94. Ribardo DA, Bingham-Ramos LK, Hendrixson DR. 2010. Functional analysis of the RdxA and RdxB nitroreductases of *Campylobacter jejuni* reveals that mutations in *rdxA* confer metronidazole resistance. J Bacteriol 192:1890–1901. doi:10.1128/JB.01638-09

95. Romilly C, Hoekzema M, Holmqvist E, Wagner EGH. 2020. Small RNAs OmrA and OmrB promote class III flagellar gene expression by inhibiting the synthesis of anti-Sigma factor FlgM. RNA Biol 17:872–880. doi:10.1080/15476286.2020.1733801

96. Ruddell B, Hassall A, Sahin O, Zhang Q, Plummer PJ, Kreuder AJ. 2020. Role of *metAB* in Methionine Metabolism and Optimal Chicken Colonization in *Campylobacter jejuni*. Infect Immun 89. doi:10.1128/IAI.00542-20

97. Scanlan E, Yu L, Maskell D, Choudhary J, Grant A. 2017. A quantitative proteomic screen of the *Campylobacter jejuni* flagellar-dependent secretome. J Proteomics 152:181–187. doi:10.1016/j.jprot.2016.11.009

98. Sharma CM, Hoffmann S, Darfeuille F, Reignier J, Findeiss S, Sittka A, Chabas S, Reiche K, Hackermüller J, Reinhardt R, Stadler PF, Vogel J. 2010. The primary transcriptome of the major human pathogen *Helicobacter pylori*. Nature 464:250–255. doi:10.1038/nature08756

99. Skouloubris S, Thiberge JM, Labigne A, De Reuse H. 1998. The *Helicobacter pylori* UreI protein is not involved in urease activity but is essential for bacterial survival in vivo. Infect Immun 66:4517–4521. doi:10.1128/IAI.66.9.4517-4521.1998

100. Stintzi A. 2003. Gene expression profile of *Campylobacter jejuni* in response to growth temperature variation. J Bacteriol 185:2009–2016. doi:10.1128/JB.185.6.2009-2016.2003

101. Sudo N, Soma A, Iyoda S, Oshima T, Ohto Y, Saito K, Sekine Y. 2018. Small RNA Esr41 inversely regulates expression of LEE and flagellar genes in enterohaemorrhagic *Escherichia coli*. Microbiology (Reading, Engl) 164:821–834. doi:10.1099/mic.0.000652

102. Sudo N, Soma A, Muto A, Iyoda S, Suh M, Kurihara N, Abe H, Tobe T, Ogura Y, Hayashi T, Kurokawa K, Ohnishi M, Sekine Y. 2014. A novel small regulatory RNA enhances cell motility in enterohemorrhagic *Escherichia coli*. J Gen Appl Microbiol 60:44–50. doi:10.2323/jgam.60.44

103. Svensson SL, Sharma CM. 2021. RNase III-mediated processing of a trans-acting bacterial sRNA and its cis-encoded antagonist. eLife 10. doi:10.7554/eLife.69064

104. Taggart JC, Lalanne J-B, Durand S, Braun F, Condon C, Li G-W. 2023. A high-resolution view of RNA endonuclease cleavage in *Bacillus subtilis*. BioRxiv. doi:10.1101/2023.03.12.532304

105. Taveirne ME, Theriot CM, Livny J, DiRita VJ. 2013. The complete *Campylobacter jejuni* transcriptome during colonization of a natural host determined by RNAseq. PLoS ONE 8:e73586. doi:10.1371/journal.pone.0073586

106. Thomason MK, Bischler T, Eisenbart SK, Förstner KU, Zhang A, Herbig A, Nieselt K, Sharma CM, Storz G. 2015. Global transcriptional start site mapping using differential RNA sequencing reveals novel antisense RNAs in *Escherichia coli*. J Bacteriol 197:18–28. doi:10.1128/JB.02096-14

107. Thomason MK, Fontaine F, De Lay N, Storz G. 2012. A small RNA that regulates motility and biofilm formation in response to changes in nutrient availability in *Escherichia coli*. Mol Microbiol 84:17–35. doi:10.1111/j.1365-2958.2012.07965.x

108. Wagner EGH, Romby P. 2015. Small RNAs in bacteria and archaea: who they are, what they do, and how they do it. Adv Genet 90:133–208. doi:10.1016/bs.adgen.2015.05.001

109. Wang Y, Taylor DE. 1990. Chloramphenicol resistance in *Campylobacter coli*: nucleotide sequence, expression, and cloning vector construction. Gene 94:23–28. doi:10.1016/0378-1119(90)90463-2

110. Waters SA, McAteer SP, Kudla G, Pang I, Deshpande NP, Amos TG, Leong KW, Wilkins MR, Strugnell R, Gally DL, Tollervey D, Tree JJ. 2017. Small RNA interactome of pathogenic *E. coli* revealed through crosslinking of RNase E. EMBO J 36:374–387. doi:10.15252/embj.201694639

111. Wei BL, Brun-Zinkernagel AM, Simecka JW, Prüss BM, Babitzke P, Romeo T. 2001. Positive regulation of motility and *flhDC* expression by the RNA-binding protein CsrA of *Escherichia coli*. Mol Microbiol 40:245–256. doi:10.1046/j.1365-2958.2001.02380.x

112. Westermann AJ. 2018. Regulatory RNAs in virulence and host-microbe interactions. Microbiol Spectr 6:RWR-0002-2017. doi:10.1128/microbiolspec.RWR-0002-2017

113. Wösten MMSM, van Dijk L, Veenendaal AKJ, de Zoete MR, Bleumink-Pluijm NMC, van Putten JPM. 2010. Temperature-dependent FlgM/FliA complex formation regulates *Campylobacter jejuni* flagella length. Mol Microbiol 75:1577–1591. doi:10.1111/j.1365-2958.2010.07079.x

114. Wösten MMSM, Wagenaar JA, van Putten JPM. 2004. The FlgS/FlgR two-component signal transduction system regulates the *fla* regulon in *Campylobacter jejuni*. J Biol Chem 279:16214–16222. doi:10.1074/jbc.M400357200

115. Yao S, Blaustein JB, Bechhofer DH. 2007. Processing of *Bacillus subtilis* small cytoplasmic RNA: evidence for an additional endonuclease cleavage site. Nucleic Acids Res 35:4464–4473. doi:10.1093/nar/gkm460

