## Supplementary Materials for "Interplay of two small RNAs fine-tunes hierarchical flagellar gene expression in the foodborne pathogen *Campylobacter jejuni*"

**This PDF file includes:**

**Supplementary figures S1-S10**

**Supplementary methods**

**Supplementary references**

### Supplementary figures

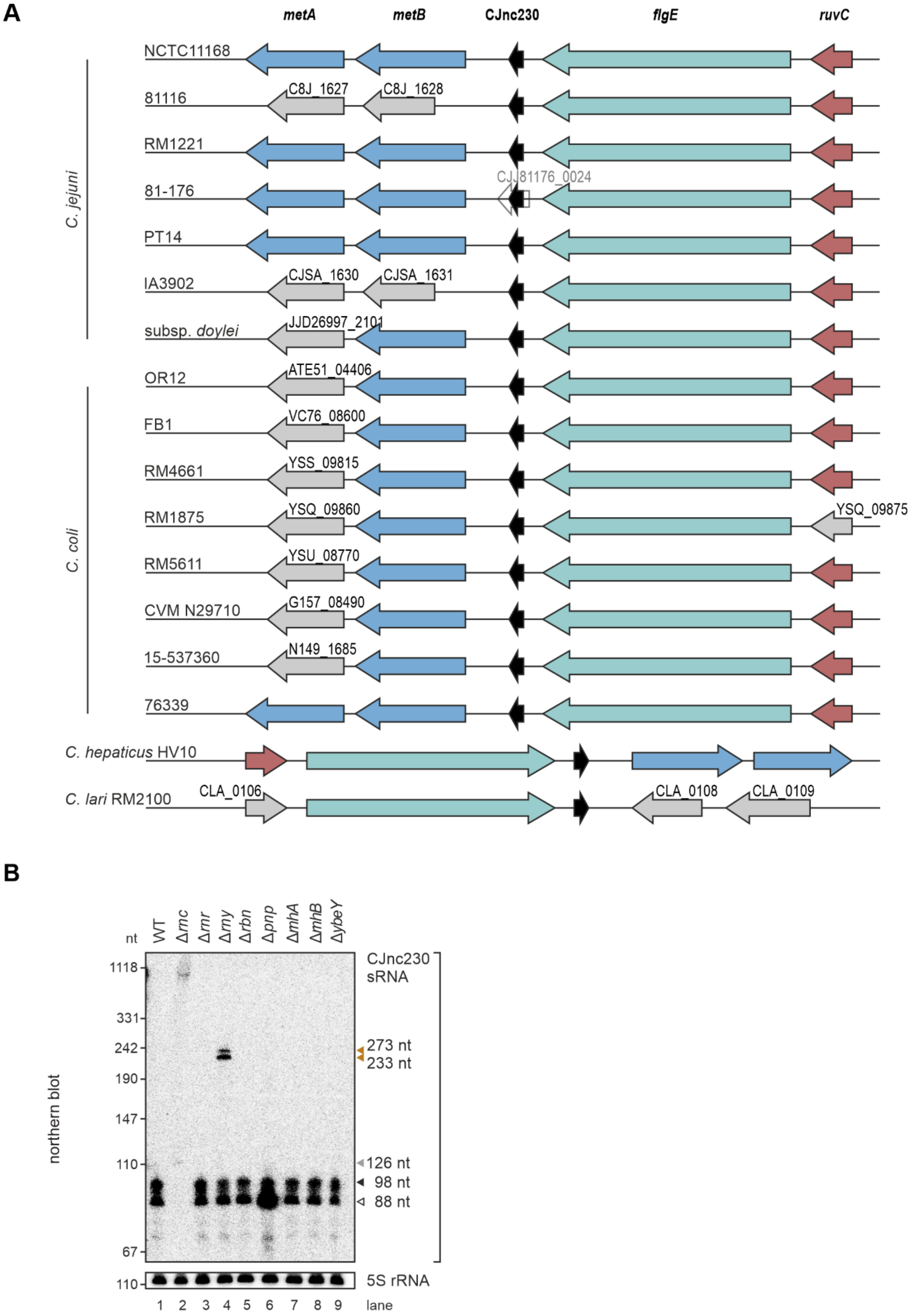


**Figure S1. CJnc230 sRNA encoded downstream of the *flgE* gene is processed by several ribonucleases. (A)** Conservational analysis of the CJnc230 genomic locus in *Campylobacter*

species and strains used for sequence alignments in **Figures 1C & S4**. Information was retrieved from KEGG (Kyoto Encyclopedia of Genes and Genomes) and orthologous genes from different strains and species are color-coded: *ruvC* - red, *flgE* (Cj1729c in NCTC11168) - turquoise, CJnc230 - black, *metB/A* - blue, hypothetical and other function - gray. For *C. jejuni* strain 81-176, a small open reading frame (CJJ81176_0024, 51 aa) is predicted to overlap with CJnc230. Sizes of arrows, indicating genes and their orientation, are not displayed at the correct scale. **(B)** Northern blot analysis of total RNA from *C. jejuni* wildtype (WT) and ribonuclease deletion strains grown to exponential phase. Transcript lengths were calculated based on 5' and 3' end positions determined by primer extension or term-seq, respectively, and marked by colored triangles (white: 88 nt, 3'-truncated version of CJnc230; black: 98 nt, full-length version of CJnc230; gray: 126 nt, possibly 5'-extended CJnc230 detected in Δ*rnc*; orange: 233/273 nt, 3'-extended versions of CJnc230 detected in Δ*rny*). Expression of CJnc230 sRNA was detected with CSO-0537 and 5S rRNA (CSO-0192) served as a loading control. Related to **Figure 2**.

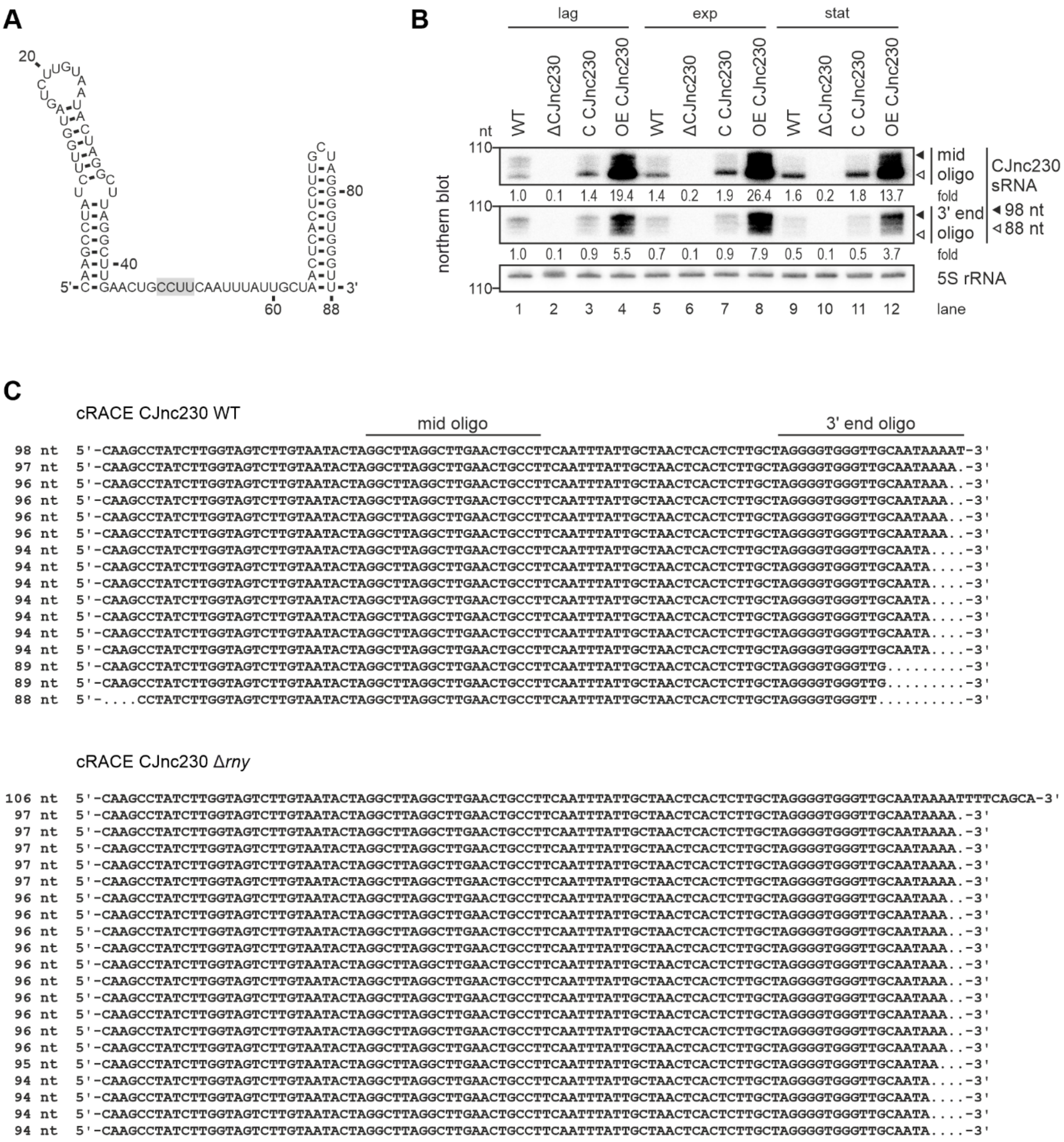


**Figure S2. The 3'-truncated version of CJnc230 increases over growth. (A)** Secondary structure of the most abundant, 3'-truncated 88-nt version of CJnc230 predicted by RNAfold (Gruber et al., 2008). Gray box: putative anti-Shine-Dalgarno motif. **(B)** Northern blot analysis of total RNA from *C. jejuni* WT and CJnc230 mutants (Δ: deletion, C: complementation in *trans*, and OE: overexpression in *trans*) harvested at lag phase (lag, $OD_{600\ nm}$ 0.1), exponential phase (exp, $OD_{600\ nm}$ ~0.4), and stationary phase (stat, $OD_{600\ nm}$ ~0.8). The full-length (black triangle, 98 nt) and the most abundant 3'-truncated version (white triangle, 88 nt) of CJnc230 are marked. CSO-0537 is complementary to the middle part of the sRNA (mid oligo), while CSO-5138 is directed towards the 3' end (3' end oligo) (positions indicated in (**C**)). 5S rRNA (CSO-0192) was used as a loading control. Fold changes of CJnc230 expression relative to WT and normalized to 5S rRNA are indicated. **(C)** Sequencing results of circular Rapid Amplification of cDNA End (cRACE) analysis of total RNA from *C. jejuni* WT and Δ*rny* strains harvested at exponential growth phase.

Depicted are CJnc230 DNA sequences with respective lengths on the left and oligonucleotide binding positions for northern blot analysis in (**B**) indicated on top.

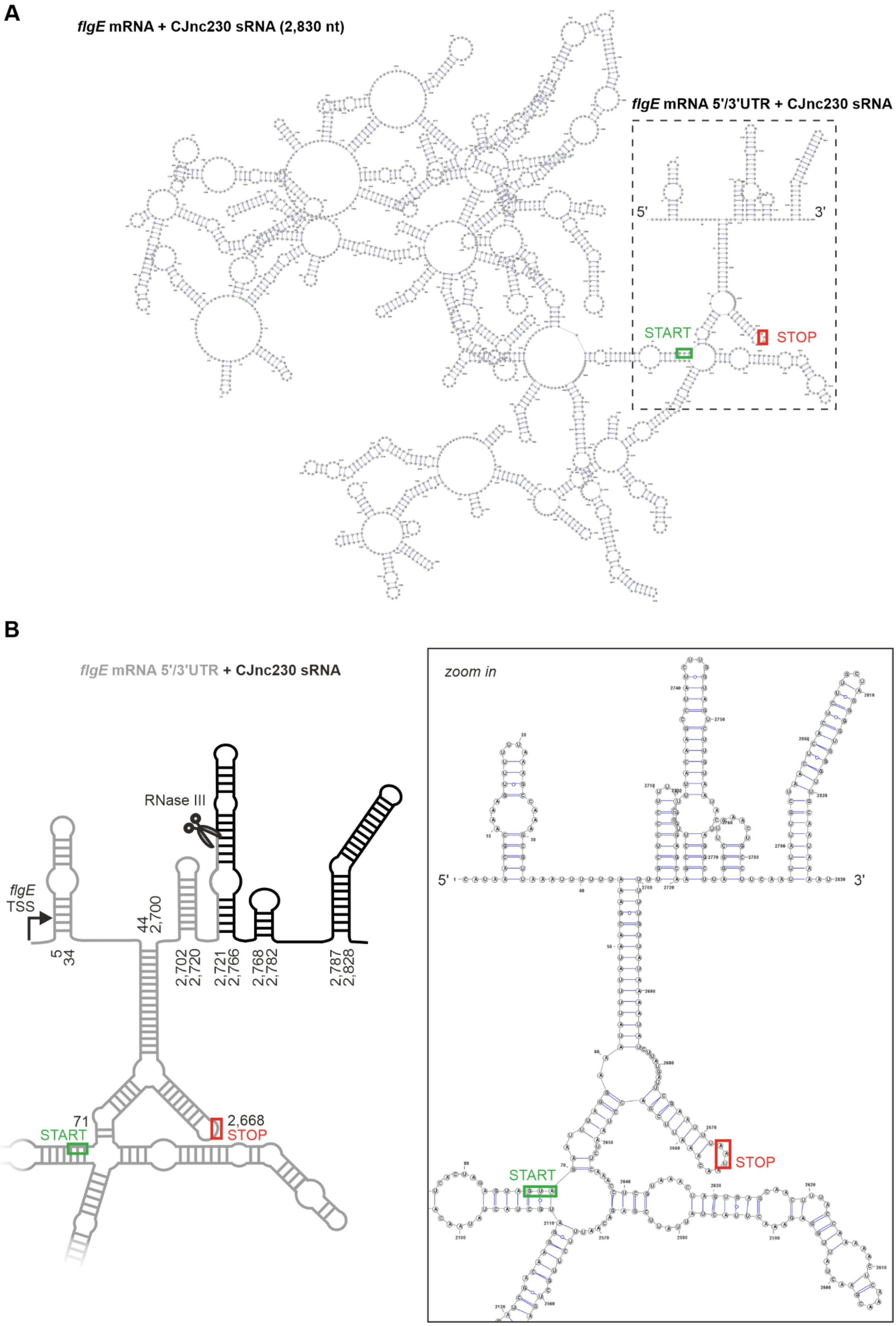
A
flgE mRNA + CJnc230 sRNA (2,830 nt)
flgE mRNA 5'/3'UTR + CJnc230 sRNA
5'
3'
START
STOP
B
flgE mRNA 5'/3'UTR + CJnc230 sRNA
RNase III
flgE
TSS
5
34
44
2,700
2,702
2,720
2,721
2,766
2,768
2,782
2,787
2,828
71
START
2,668
STOP
zoom in
5'
3'
START
STOP

**Figure S3. Secondary structure prediction reveals a stem-loop for possible RNase III cleavage downstream of the *flgE* CDS. (A)** RNA secondary structure prediction of the 2,830-nt long *flgE*-CJnc230 transcript of *C. jejuni* NCTC11168 by RNAfold (Gruber et al., 2008). The dashed box highlights the *flgE* (Cj1729c) start (green box) and stop (red box) codon, 5'UTR and 3'UTR including the CJnc230 sRNA, and transcript 5' and 3' ends illustrated in more detail in (**B**). **(B)** Scheme (*left*) and corresponding zoom in region (*right*) of potential RNase III cleavage (scissors) in a stem-loop downstream of the *flgE* CDS, thus producing the CJnc230 sRNA (black) 5' end. Bent arrow: transcriptional start site (TSS) of the *flgE* mRNA (Porcelli et al., 2013). Stem-loop positions with respect to the TSS are provided in the scheme to estimate sizes. Start (green box) and stop (red box) codon positions of the *flgE* open reading frame are also indicated.

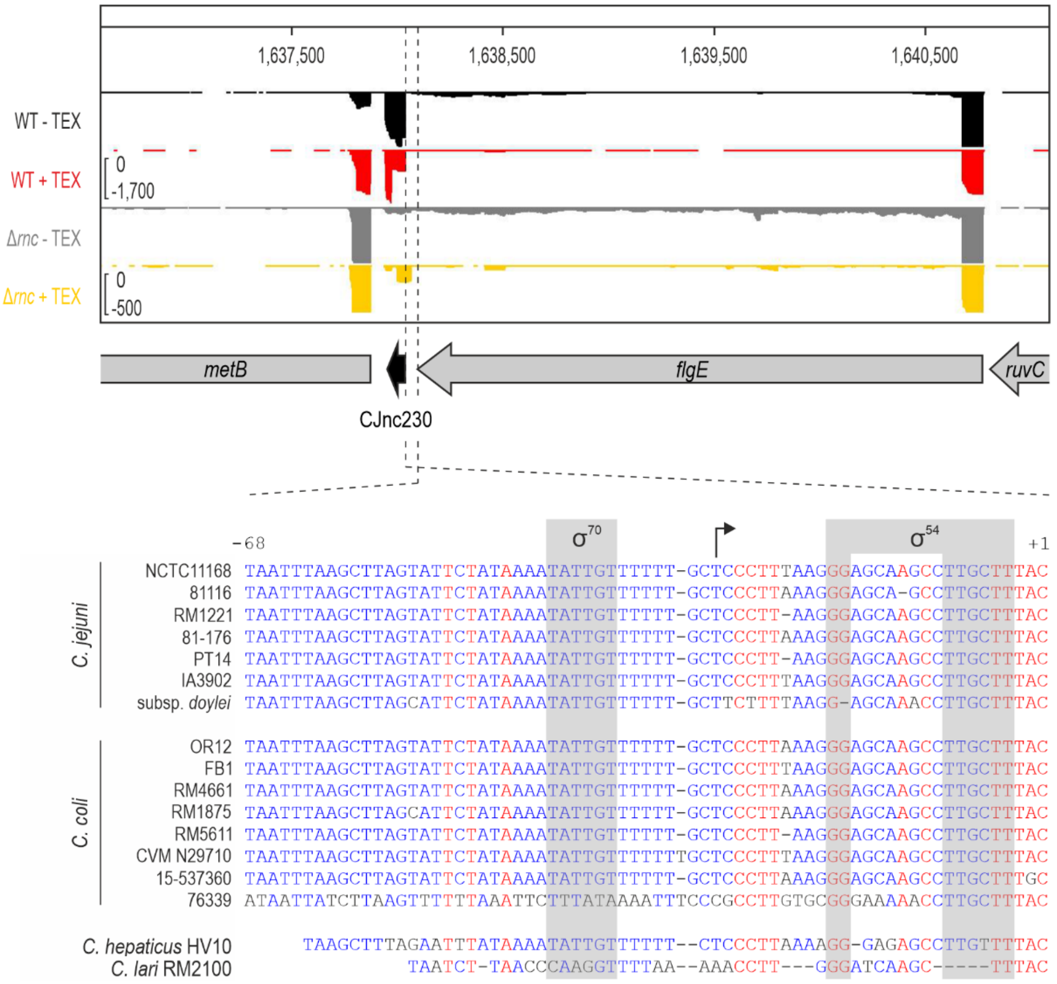


**Figure S4. Differential RNA sequencing (dRNA-seq) reveals an alternative TSS upstream of CJnc230.** (*Upper*) dRNA-seq coverage of the *flgE*-CJnc230 locus in *C. jejuni* WT and the Δ*rnc* mutant grown to exponential phase. -/+TEX: mock-/terminator exonuclease (TEX)-treated dRNA-seq (Sharma et al., 2010) libraries. TEX enriches for 5'-triphosphorylated primary transcript ends, as a consequence of degrading processed (non-triphosphorylated) 5' ends. (*Lower*) Genomic sequence alignment by MultAlin (Corpet, 1988) of multiple *Campylobacter* species and strains comprising the *flgE* 3'UTR from the stop codon (TAA, positions -68 to -66) to the CJnc230 5' end (processing site, C residue, +1). Gray boxes: two promoter motifs for $\sigma^{70}$ and $\sigma^{54}$, respectively. Bent arrow: TSS detected by dRNA-seq and primer extension (**Fig. 2B**) in Δ*rnc*.

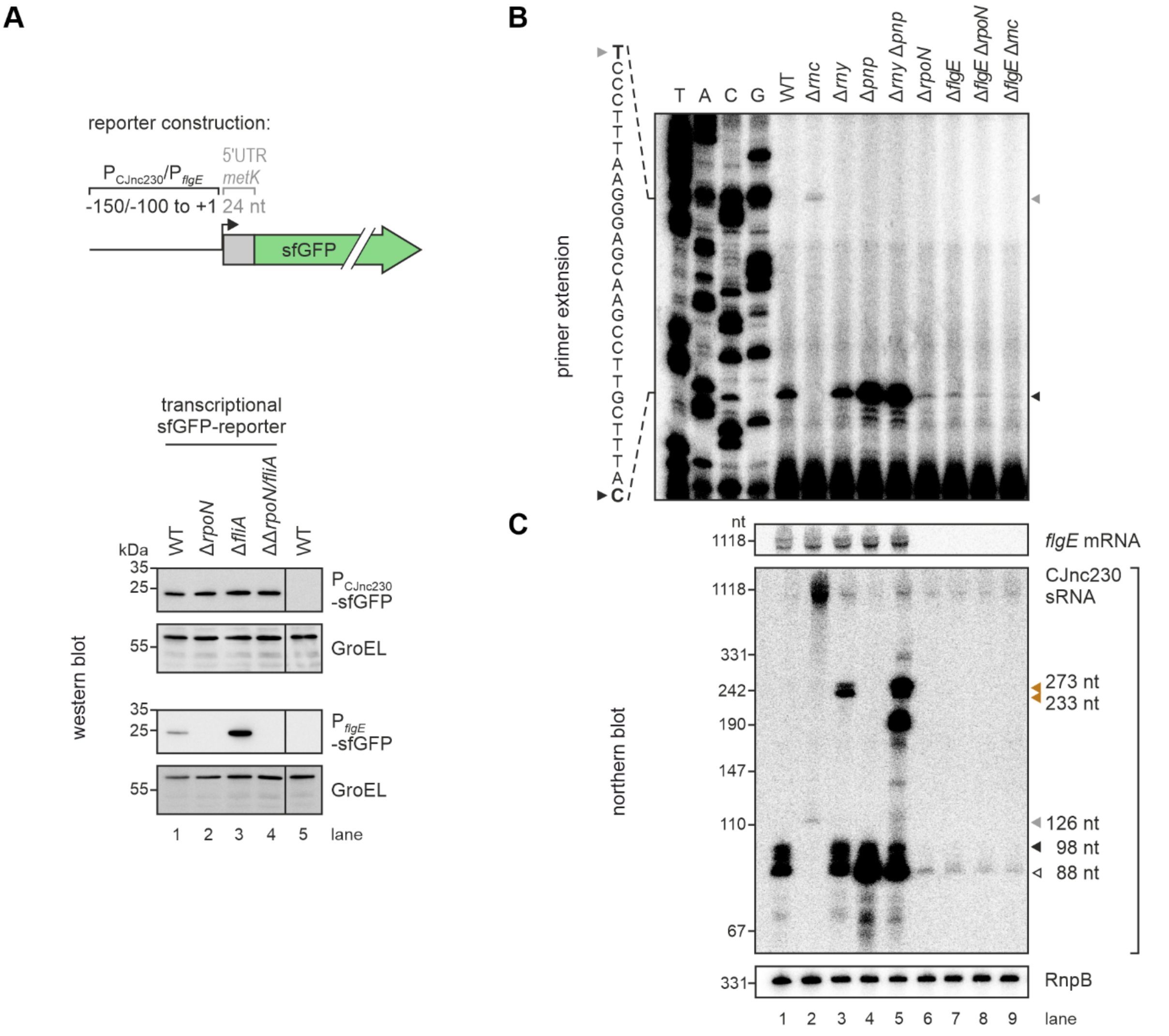


**Figure S5. A minor fraction of CJnc230 sRNA originates from an RpoD-dependent promoter. (A)** (*Upper*) *C. jejuni* transcriptional reporter construction. Either 100 or 150 nucleotides upstream of the *flgE* TSS (Porcelli et al., 2013) or the CJnc230 5' end/processing site (**Fig. 2B**) were fused to an unrelated ribosome binding site (RBS) (*metK*) and *sfgfp*. This construct was introduced together with a resistance cassette into the Cj0046 pseudogene locus. (*Lower*) Western blot analysis of $P_{CJnc230}$ and $P_{flgE}$ transcriptional reporters in *C. jejuni* WT and sigma factor deletion strains grown to exponential phase. GroEL was detected for normalization. Representative images were cut between lanes 4 and 5. **(B)** Primer extension analysis of total RNA from *C. jejuni* WT, ribonuclease deletion mutants, and Δ*rpoN*, Δ*flgE*, or Δ*flgE* mutants combined with deletion of *rpoN* or *rnc*, harvested at exponential phase. Lanes 1-5 are identical to main **Figure 2B**. Total RNA was annealed with the probe for CJnc230 used for northern blots (CSO-0537), binding in the middle of the sRNA. A sequencing ladder generated with this probe is partially indicated on the left. The gray triangle marks the alternative RpoD-dependent TSS (T residue in bold on the left) detected in Δ*rnc* bacteria (**Fig. S4**), 28 nt upstream of the sRNA 5' end (C residue in bold on the left, black triangle) in WT. **(C)** Northern blot analysis of total RNA from *C. jejuni* WT, ribonuclease deletion mutants, and Δ*rpoN*, Δ*flgE*, or Δ*flgE* mutants combined with deletion of *rpoN* or *rnc*, harvested at exponential growth phase. Lanes 1-5 are identical to main **Figure 2A**. Lengths of prominent CJnc230 transcripts are indicated by colored triangles and were determined by term-seq and/or primer extension. Expression of CJnc230 sRNA was detected with CSO-0537, *flgE* mRNA with CSO-5136 (binding the *flgE* CDS), and RnpB RNA (CSO-0497) served as a loading control.

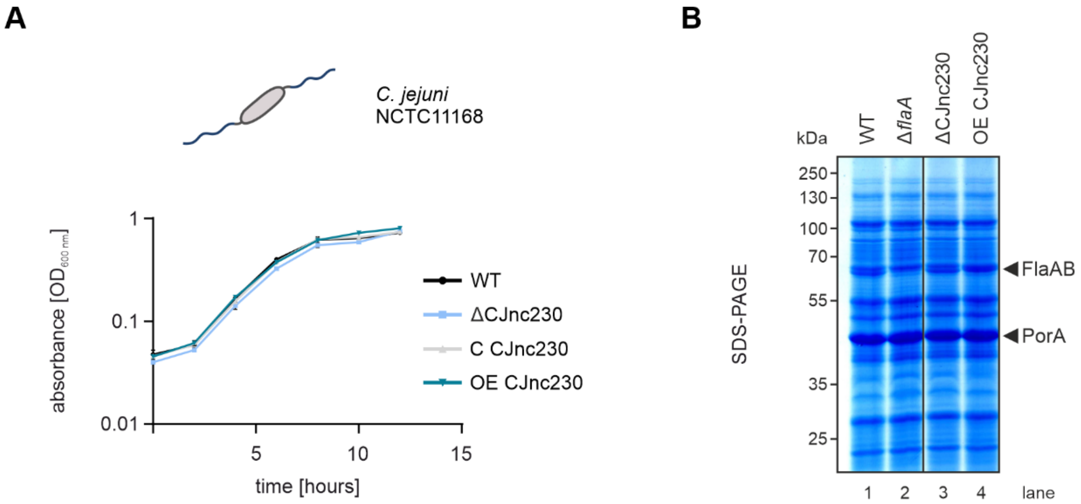


**Figure S6. Growth behavior and analysis of total protein patterns of *C. jejuni* CJnc230 mutant strains. (A)** Growth curve analysis of *C. jejuni* WT and CJnc230 deletion (Δ), complementation (C), and overexpression (OE) mutants. Strains were grown for 12 hours in Brucella broth (BB) medium and culture density ($OD_{600\ nm}$) was measured every 2 hours. The mean of two independent replicates ± standard deviation is shown. **(B)** Total protein samples from *C. jejuni* WT and CJnc230 mutant strains (Δ and OE) harvested at exponential phase. On the SDS-PAGE image, the flagellins (FlaA/B) and the non-regulated major outer membrane protein PorA (loading control) are indicated. Δ*flaA*: negative control. Image was cut between lanes 2 and 3.

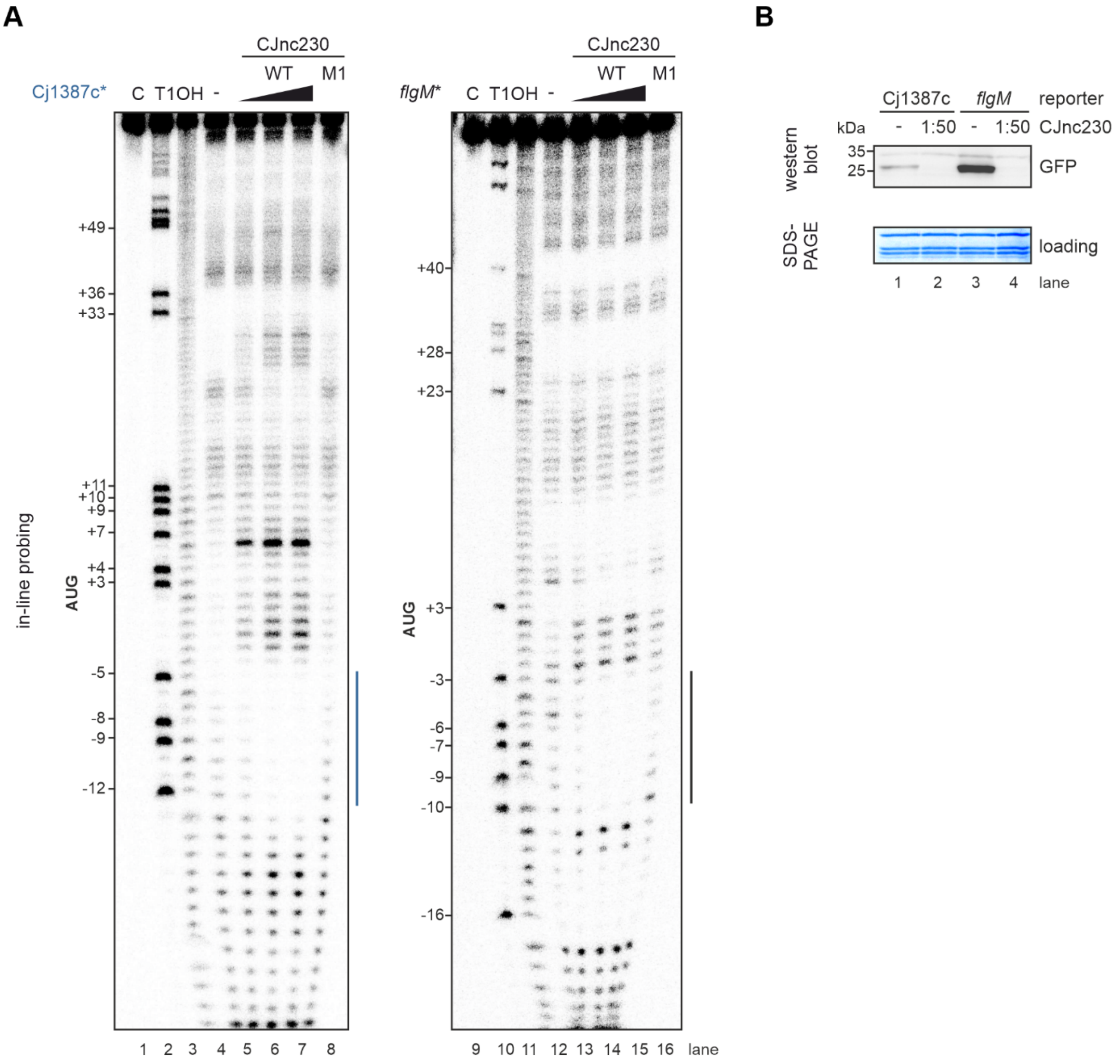


**Figure S7. *In-vitro* in-line probing and translational reporter assays confirm Cj1387c and *flgM* mRNAs as direct targets of CJnc230 repression. (A)** In-line probing of 0.2 pmol $^{32}$P-5'-end-labeled (marked with *) Cj1387c and *flgM* mRNA leaders in the absence or presence of 0.02/0.2/2 pmol unlabeled CJnc230 WT sRNA or 2 pmol M1 mutant sRNA. Interaction sites with Cj1387c (blue) and *flgM* (black) 5'UTRs are indicated on the right. C - untreated control; T1 ladder - G residues (indicated on the left); OH - all positions (alkaline hydrolysis). **(B)** *In-vitro* translation of Cj1387c- and *flgM*-sfGFP reporters (5'UTR and first 10 codons fused to *sfgfp*, 4 pmol) in an *E. coli* cell-free system -/+ CJnc230 (1:50 = 200 pmol) detected by western blot. PageBlue staining of the gel after blotting served as a loading control.

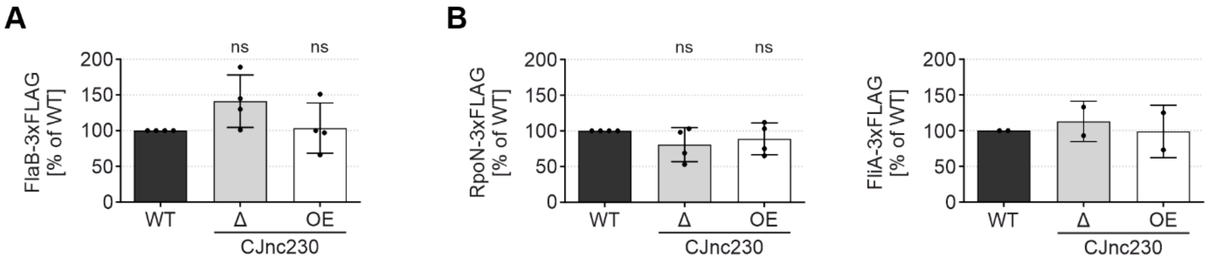


**Figure S8. CJnc230 does not affect levels of the minor flagellin FlaB or flagellar sigma factors. (A and B)** Protein levels of C-terminally FLAG-tagged FlaB **(A)** or RpoN (*left*) and FliA (*right*) **(B)** in *C. jejuni* grown to exponential phase measured by western blot. Bar graphs represent the mean of independent replicates (n = 4 for FlaB-3xFLAG; n = 4 for RpoN-3xFLAG; n = 2 for FliA-3xFLAG), error bars depict the standard deviation. ns: not significant, two-tailed Student's *t*-test was used to compare the respective mutant to WT.

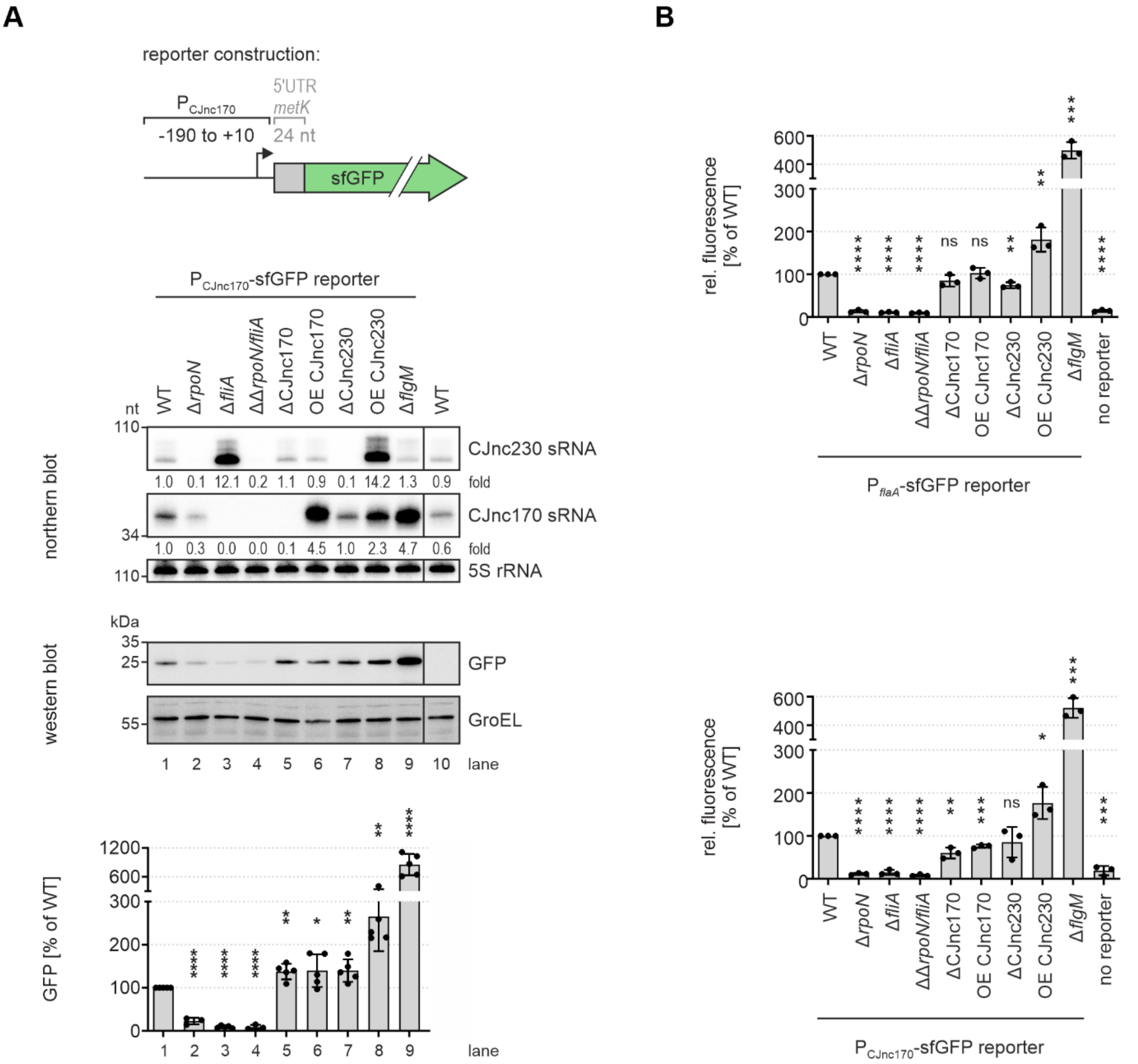


**Figure S9. Transcriptional reporter assays confirm CJnc230 activation of class III flagellar gene transcription. (A)** Northern and western blot analyses of *C. jejuni* WT or sigma factor, sRNA, and anti-sigma factor mutant strains with a transcriptional reporter of the $P_{CJnc170}$ promoter region fused to an unrelated RBS (*metK*) and *sfgfp*, harvested at exponential growth phase. Nucleotide positions with respect to the CJnc170 TSS (Dugar et al., 2013; Porcelli et al., 2013) are indicated in the scheme on top. (*Middle panel*) Northern blot validation of CJnc230 (CSO-0537) and CJnc170 (CSO-0182) sRNA expression. 5S rRNA (CSO-0192) was used as a loading control. Fold changes of sRNA expression relative to the WT reporter and normalized to 5S rRNA are indicated. Images were cut between lanes 9 and 10. (*Lower panel*) Reporter expression in the respective mutants was measured by western blotting. GroEL was detected for normalization. One representative blot (cut between lanes 9 and 10) is shown and quantification of independent replicates (n = 5 for WT, Δ*fliA*, ΔCJnc170, OE CJnc170, ΔCJnc230, OE CJnc230, and Δ*flgM*; n = 3 for Δ*rpoN* and Δ*rpoN*/Δ*fliA*) is depicted in the bar graph below. Error bars represent the standard deviation. ****: $p < 0.0001$, **: $p < 0.01$, *: $p < 0.05$, ns: not significant, two-tailed Student's *t*-test was used to compare the respective mutant to the WT reporter background (lane 1). **(B)** Flow cytometry analyses of $P_{flaA}$ (*upper*) and $P_{CJnc170}$ (*lower*) transcriptional reporter expression at exponential growth phase. Bar graphs based on 100,000 counted cells per sample represent the mean of independent replicates (n = 3), error bars depict the standard deviation. ****: $p < 0.0001$, ***: $p < 0.001$, **: $p < 0.01$, *: $p < 0.05$, ns: not significant, two-tailed Student's *t*-test was used to compare the respective mutant to the WT reporter background (WT).

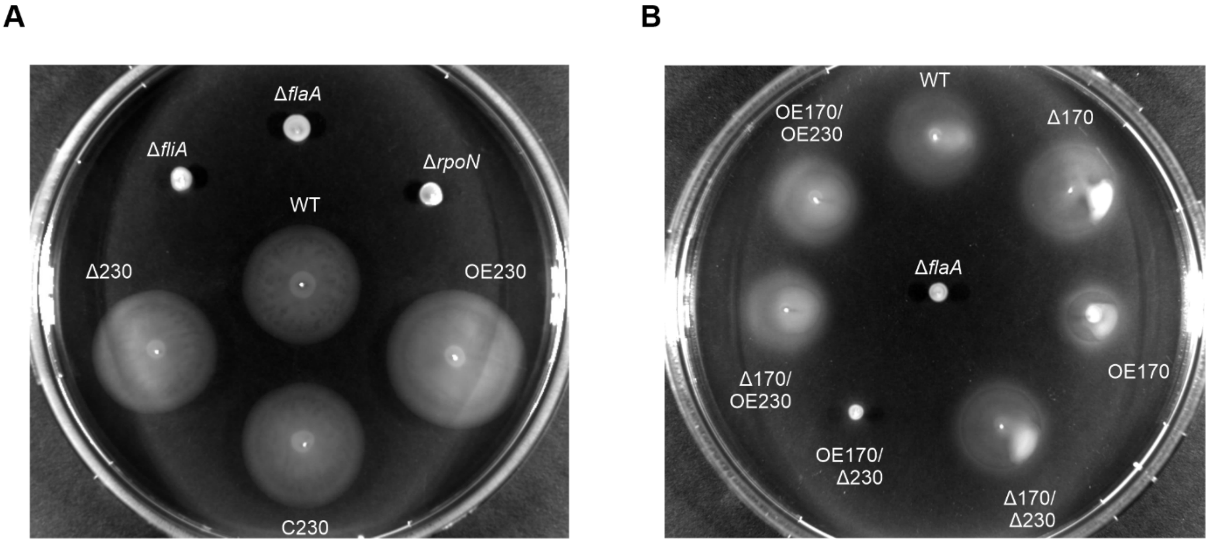


**Figure S10. Motility assays with sRNA single or double mutants reveal opposite phenotypes and motility regulation.** Representative images of swimming motility assays of *C. jejuni* WT and CJnc230 mutant strains (Δ230: CJnc230 deletion, C230: CJnc230 complementation in *trans*, OE230: CJnc230 overexpression in *trans*) **(A)** or CJnc170 deletion (Δ170) or overexpression (OE170) mutants with or without deletion or overexpression of CJnc230 **(B)** in 0.4% soft agar BB plates. Δ*flaA*/Δ*fliA*/Δ*rpoN*: non-motile controls lacking major flagellin or flagellar sigma factors FliA and RpoN. Related to main **Figures 4B & 6A**.

### Supplementary methods

#### Circular Rapid Amplification of cDNA Ends (cRACE).

To simultaneously map CJnc230 transcript 5' and 3' ends in *C. jejuni* NCTC11168 wildtype (WT) and the Δ*rny* mutant, Rapid Amplification of cDNA Ends (RACE) after self-ligation of RNA transcripts was performed.

First, 5 µg of DNase I-digested RNA was taken up in a total volume of 16.5 µl $H_2O$, denatured, and snap-cooled on ice. A mix of 2 µl 10 x Antarctic Phosphatase buffer (NEB), 1 µl Antarctic Phosphatase (5 U; NEB), and 0.5 µl Ribonuclease Inhibitor (10 U; Molox) was added and incubated for 1 h at 37°C to dephosphorylate the RNA. The reaction was then made up to 100 µl and extracted with an equal volume of phenol:chloroform:isoamyl (PCI) alcohol in a Phase-Lock gel tube (5PRIME). Afterwards, RNA was precipitated with 1 µl GlycoBlue™ (Thermo Fisher Scientific) and 2.5 volumes 30:1 mix (100% EtOH:3 M sodium acetate, pH 6.5). The RNA was dissolved in 11.5 µl $H_2O$ and 8.5 µl of a mix containing 2 µl 10 x T4 RNA ligase buffer (NEB), 3 µl DMSO, 2 µl ATP (1 mM final concentration; NEB), 1 µl T4 RNA ligase 1 (10 U; NEB), and 0.5 µl Ribonuclease Inhibitor (10 U; Molox) was added. Following ligation at 37°C for 30 min, RNA was extracted with PCI and precipitated as described above. Then, it was solved in 30 µl $H_2O$ and 10 µL of circularized RNA, together with 2 µl dNTPs (1 mM final concentration) and 1 µl reverse transcription (RT) primer (CSO-0537; 5 µM final concentration), were denatured and snap-cooled on ice. The RT reaction was performed with 4 µl 5 x RT buffer (Thermo Scientific), 1 µl Ribonuclease Inhibitor (20 U; Molox), 1 µl 0.1 M DTT, and 1 µL Maxima RT (200 U; Thermo Scientific) for 5 min at 50°C, 1 h at 55°C, and 15 min at 70°C. RNA was removed by digestion with 1 µl RNase H (5 U; NEB) for 22 min at 37°C.

One microliter of this reaction was used as a template for PCR using *Taq* DNA polymerase (NEB) with oligonucleotides CSO-0537 x 5192 and 3% DMSO. Cycling conditions were as follows: 95°C for 5 min and 35 cycles of [95°C, 10 s; 56°C, 30 s; 72°C, 60 s], followed by 72°C for 10 min. Amplification was checked on an 8% PAA gel, reactions were cleaned up with the NucleoSpin Gel and PCR Clean-up kit (Macherey-Nagel), and ligated into pGEM-T Easy (Promega) according to the manufacturer's instructions. For the WT background, the inserts of sixteen white clones were sequenced, and for Δ*rny* bacteria, the inserts of twenty-one white clones were sequenced with primers REV or UNI-61.

**Differential RNA sequencing (dRNA-seq).**

For comparative TSS analysis in *C. jejuni* NCTC11168 WT and RNase III-deficient bacteria, bacterial total RNA samples were harvested at mid-log phase after growth in rich BB medium. After digestion of residual genomic DNA (gDNA) by DNase I treatment, the sample was equally divided into two parts and Terminator 5'-phosphate-dependent exonuclease (TEX) (Epicentre) was used to deplete processed transcripts in one half as previously described (Sharma et al., 2010). Libraries for Solexa sequencing (HiSeq) were constructed at Vertis Biotechnologie AG, Germany, as previously described (Berezikov et al., 2006), but without RNA fractionation prior to cDNA synthesis. In brief, equal amounts of RNA were poly(A)-tailed using poly(A) polymerase and 5'-triphosphate groups were digested with tobacco acid pyrophosphatase (TAP). Then, an RNA adapter was ligated to the 5' phosphate and first-strand cDNA synthesized with an oligo(dT)-adapter primer and M-MLV (Moloney Murine Leukaemia Virus) reverse transcriptase. A high-fidelity DNA polymerase was used to PCR-amplify the resulting cDNA to a concentration of ~20-30 ng/µl. Finally, cDNA was purified with the Agencourt AMPure XP kit (Beckman Coulter Genomics), analyzed by capillary electrophoresis, and sequenced on an Illumina HiSeq 2000 platform in single-end mode.

The FASTQ-format reads were trimmed with a cutoff Phred score of 20 using Cutadapt (version: 4.1) (Martin, 2011). After poly(A)-tails were removed, sequences shorter than 12 nt were omitted and the remaining reads mapped to the *C. jejuni* NCTC11168 reference genome (NCBI accession number: NC_002163.1, RefSeq assembly accession number: GCF_000009085.1) using READemption (version: 2.0.1) (Förstner et al., 2014) and segemehl (version: 0.3.4) (Hoffmann et al., 2009) with an accuracy cutoff of 95%. Coverage plots containing the number of mapped reads per nucleotide were also generated with READemption and visualized in the Integrated Genome Browser (Freese et al., 2016). The coverage was normalized by the total number of aligned reads of a given library and multiplied by the minimum number of aligned reads calculated over all libraries. Gene annotations were retrieved from NCBI and extended as described for total RNA sequencing experiments.

### Supplementary references


Berezikov E, Thuemmler F, van Laake LW, Kondova I, Bontrop R, Cuppen E, Plasterk RHA. 2006. Diversity of microRNAs in human and chimpanzee brain. *Nat Genet* **38**:1375–1377. doi:10.1038/ng1914

Corpet F. 1988. Multiple sequence alignment with hierarchical clustering. *Nucleic Acids Res* **16**:10881–10890. doi:10.1093/nar/16.22.10881

Dugar G, Herbig A, Förstner KU, Heidrich N, Reinhardt R, Nieselt K, Sharma CM. 2013. High-resolution transcriptome maps reveal strain-specific regulatory features of multiple *Campylobacter jejuni* isolates. *PLoS Genet* **9**:e1003495. doi:10.1371/journal.pgen.1003495

Förstner KU, Vogel J, Sharma CM. 2014. READemption-a tool for the computational analysis of deep-sequencing-based transcriptome data. *Bioinformatics* **30**:3421–3423. doi:10.1093/bioinformatics/btu533

Freese NH, Norris DC, Loraine AE. 2016. Integrated genome browser: visual analytics platform for genomics. *Bioinformatics* **32**:2089–2095. doi:10.1093/bioinformatics/btw069

Gruber AR, Lorenz R, Bernhart SH, Neuböck R, Hofacker IL. 2008. The Vienna RNA websuite. *Nucleic Acids Res* **36**:W70-4. doi:10.1093/nar/gkn188

Hoffmann S, Otto C, Kurtz S, Sharma CM, Khaitovich P, Vogel J, Stadler PF, Hackermüller J. 2009. Fast mapping of short sequences with mismatches, insertions and deletions using index structures. *PLoS Comput Biol* **5**:e1000502. doi:10.1371/journal.pcbi.1000502

Martin M. 2011. Cutadapt removes adapter sequences from high-throughput sequencing reads. *EMBnet j* **17**:10. doi:10.14806/ej.17.1.200

Porcelli I, Reuter M, Pearson BM, Wilhelm T, van Vliet AHM. 2013. Parallel evolution of genome structure and transcriptional landscape in the Epsilonproteobacteria. *BMC Genomics* **14**:616. doi:10.1186/1471-2164-14-616

Sharma CM, Hoffmann S, Darfeuille F, Reignier J, Findeiss S, Sittka A, Chabas S, Reiche K, Hackermüller J, Reinhardt R, Stadler PF, Vogel J. 2010. The primary transcriptome of the major human pathogen *Helicobacter pylori*. *Nature* **464**:250–255. doi:10.1038/nature08756
